# Extremely potent Human Monoclonal Antibodies for the Prophylaxis and Therapy of Tetanus

**DOI:** 10.1101/2021.05.24.445390

**Authors:** Marco Pirazzini, Alessandro Grinzato, Davide Corti, Sonia Barbieri, Oneda Leka, Francesca Vallese, Marika Tonellato, Chiara Silacci-Fregni, Luca Piccoli, Eaazhisai Kandiah, Giampietro Schiavo, Giuseppe Zanotti, Antonio Lanzavecchia, Cesare Montecucco

**Author notes:** These authors contributed equally (Marco Pirazzini, Alessandro Grinzato).

## Abstract

Human monoclonal antibodies were used here to study the mechanism of neuron intoxication by tetanus neurotoxin protein toxins and as a safe preventive and therapeutic substitute of hyperimmune sera. By screening memory B cells of immune donors, we selected two monoclonal antibodies specific for tetanus neurotoxin with exceptionally high neutralizing activities, which have been extensively characterized both structurally and functionally. We found that these antibodies interfere with the binding and translocation of the neurotoxin into neurons by interacting with two epitopes, whose definition pinpoints crucial events in the cellular pathogenesis of tetanus. Some mechanistic aspects of tetanus neurotoxin intoxication were revealed, explaining at the same time, the unprecedented neutralization ability of these antibodies. Importantly, these antibodies are exceptionally efficient in preventing experimental tetanus when injected in mice long before the neurotoxin. Moreover, their Fab derivatives neutralize tetanus neurotoxin in post-exposure experiments, suggesting their potential therapeutic use upon intrathecal injection. As such, these human monoclonal antibodies, as well as their Fab derivatives, meet all requirements for being considered for prophylaxis and therapy of human tetanus and are ready for clinical trials.

## INTRODUCTION

Tetanus neurotoxin (TeNT) is a highly potent exotoxin responsible for tetanus, a life-threatening disease whose major symptoms are muscle rigidity and spasms, spastic paralysis, respiratory deficits and autonomic dysfunctions (1-5). TeNT is produced by toxigenic strains of the anaerobic sporogenic bacterium *Clostridium tetani* (6-8). The few amino acid variations found in the currently known TeNT isoforms do not change their immunogenic properties with respect to the prototypical TeNT (Harvard strain, E88) (9).

TeNT consists of a metalloprotease light chain (L, 50 kDa) and a heavy chain (H, 100 kDa) linked by a single interchain disulfide bond essential for neurotoxicity (8, 10). TeNT is generally described as consisting of three domains, each serving a different step of its mechanism of neuronal intoxication. The carboxy-terminal domain HC (50 kDa) is responsible for presynaptic binding and consists of two sub-domains; the carboxy-terminal half (HC-C) contains a polysialoganglioside binding site and a nidogen binding site (11-14), whereas the amino-terminal half portion (HC-N), is essential for toxicity though its function is not known (15). HC-N is linked to HN (50 kDa), the domain responsible for the membrane translocation of the L domain into the cytosol. The L domain is a metalloprotease that causes tetanus by blocking the release of neurotransmitters from inhibitory interneurons of the spinal cord that control the balanced contraction of efferent motor neurons (2, 5, 16).

*C. tetani* spores are ubiquitous in the environment and can contaminate necrotic wounds of any kind, including burns, ulcers, surgery, tattooing, circumcision and needle injection. Inside these lesions, the spores may generate vegetative bacteria producing TeNT which diffuses via the blood and lymphatic circulations. It binds to motor, sensory and autonomic presynaptic nerve terminals via at least two independent receptors: a polysialoganglioside and a protein (5, 17). Polysialogangliosides (PSG), including GT1b, GD1b and GQ1b, play a major role in the initial membrane binding of TeNT (11-13), whilst nidogens 1 and 2 (also known as entactin 1/2) were identified as TeNT protein receptors (14). Nidogens and PSG direct TeNT to presynaptic zones that, upon endocytosis, generate signaling endosomes containing neurotrophic factors and their receptors (18). These TeNT-positive organelles undergo fast axonal retrograde transport from the periphery to the perikaryon located in the spinal cord, where they release their content (18-20). TeNT then binds to inhibitory interneurons and is endocytosed into the lumen of synaptic vesicles (SV) (21). The protein SV2 has been suggested to act as the protein receptor involved in this process (22, 23). Acidification of the SV lumen induces a change in conformation of TeNT whereby the HN domain inserts into the membrane and assists the membrane translocation of the L chain from the SV lumen to the cytosol. Here, the interchain disulfide bond is reduced by the thioredoxin reductase-thioredoxin redox system (24), releasing the metalloprotease activity of the L chain, which specifically cleaves VAMP-1/2/3 at a single site (25). Together with SNAP-25 and syntaxin, VAMP forms a complex driving the fusion of SV with the presynaptic membrane with ensuing neurotransmitter release (26).

VAMP cleavage in spinal cord inhibitory interneurons prevents the release of inhibitory neurotransmitters, which determines the hyperactivation of the post-synaptic motor neurons with sustained contraction of the innervated skeletal muscles resulting in spastic paralysis (27, 28). Muscle contractures begin in the head with *trismus* (lockjaw, a cardinal symptom of tetanus) and neck with difficult swallowing and then descend to the thorax, causing respiratory deficit, generalized spasm of opposing skeletal muscles and autonomic dysfunctions that may lead to death (2).

Tetanus is prevented by a very effective vaccine based on tetanus toxoid consisting of a formaldehyde-inactivated TeNT. Repeated injections of this vaccine in childhood raise a long-lasting protection due to specific anti-TeNT antibodies produced by long-lived plasma cells. After the age of forty/fifty, boost injections at 10-year intervals is suggested (CDC, 2006; WHO, 2017), in view of slowly decreasing antibody titers (29). Anti-tetanus vaccination does not generate herd immunity that protects non immunized individuals because TeNT is not infectious. Despite this, tetanus is still a major killer in countries where appropriate public healthcare systems are not enforced, and where the infection of the umbilical cord stump or of the birth canal with non-sterile instruments causes *tetanus neonatorum* and/or maternal tetanus. Neonatal tetanus frequently leads to a horrible death within a few days; maternal tetanus is life-threatening as well (30).

Another common medical practice to prevent the development of tetanus in patients presenting wounds at hospital emergency rooms is the passive immunization generated by the injection of anti-TeNT IgG, known as tetanus immunoglobulins (TIG), isolated from hyperimmune human donors (31). Hyperimmune horse sera are used in poor countries because of their lower cost but can generate dangerous hypersensitivity reactions. TIG is also administered to patients with active tetanus, to neutralize circulating TeNT in body fluids and limit the severity of the disease (31). Intrathecal TIG administration is more effective, although this approach is limited by the low percentage of anti-TeNT antibodies present in TIG and the total amount of protein that can be injected (32, 33).

The hyperimmune human sera used in the past exposed the patients to a risk of accidental infection with viruses, such as human HIV, hepatitis and parvovirus B19, but this possibility has been nowadays reduced by selection of donors and better protocols of IgG isolation. Yet the possibility of contamination with unknown pathogens present in the original blood samples cannot be excluded *a priori*. Moreover, the vast majority of TIG IgGs are not specific for TeNT. As a result, the usual 250 - 500 I.U. immunoglobulin injected dose has a high protein content (typically 16 – 20 % w/v), which may elicit adverse reactions such as angioedema or anaphylaxis. In addition, the anti-tetanus human immunoglobulins currently used contain IgA and this may cause in an immune reaction in IgA-deficient subjects. Pain at the site of injection or fever, dizziness and other side effects are not uncommon. An additional drawback of TIG is the intrinsic lot-to-lot variability. These drawbacks could be overcome by using highly purified human monoclonal antibodies (humAbs), which are currently used for treating a variety of human diseases (34). Methods to immortalize pathogen specific human B memory cells have been optimized (35, 36) and used to isolate human monoclonal antibodies endowed with high neutralizing properties against a variety of pathogens (37-40).

Here, we report on the properties of a TeNT-specific repertoire of humAbs and their characterization in terms of binding to the different structural domains of TeNT. We identified two humAbs endowed with an extremely high neutralizing activity that recognize the HC-C domain and the HN domain TeNT. We generated the corresponding Fab and isolated a ternary TeNT immunocomplex that was analyzed by cryo-electron microscopy. The molecular structure of this complex explains the high neutralizing power of the two antibodies at nearly stoichiometric ratio and identifies the specific steps of the process of TeNT entry into neurons blocked by these antibodies. In addition, these humAbs provided a long-lasting prophylactic activity against lethal doses of TeNT whereas their Fab prevented tetanus in post-exposure experiments as well as TIG. Thus, these findings suggest that these humAbs and their Fab fragments represent appropriate and safe alternatives to TIG.

## RESULTS

### HumAbs recognize distinct domains of TeNT and display distinct neutralizing capability

Human monoclonal antibodies against TeNT were isolated from memory B cells of an immune donor that underwent routine vaccinations with tetanus toxoid as described (35). IgG memory B cells were immortalized under clonal conditions and the antibodies produced in the culture supernatant were screened by ELISA for their ability to bind TeNT. The antibody genes from 14 positive clonal cultures were sequenced and recombinant antibodies were produced as IgG1 by transfecting HEK-293 cells.

The TeNT domains recognized by the fourteen humAbs were determined by western blotting of the intact or reduced toxin and of the recombinant sub-domains (Figure 1). Data documenting the binding specificity of each antibody are reported in Figure S1. Most antibodies bound to the HC domain, three to the HN domain and none to the L chain. While three of the eleven HC-specific antibodies recognized the C-terminal HC-C part, eight bound to the HC-N portion (Figure S1), suggesting that HC-N is the most immunogenic part of TeNT. The specific binding to HC-N and/or HC-C was not investigated in recent studies on anti-TeNT humAbs, which reported that antibodies binding to intact HC (HC-N plus HC-C) displayed the highest neutralizing power (41, 42). Although the role of HC-N in neuron intoxication is currently unknown, its deletion causes loss of toxicity (15). Therefore, these specific humAbs could be very useful in unravelling the role of HC-N in the molecular and cellular pathogenesis of tetanus. Together with the knowledge that HC-C contains the binding sites for the neuronal receptors of TeNT, these findings strongly suggest that the structure of TeNT is best described in terms of four domains (L, HN, HC-N and HC-C), rather than the three-domain paradigm reported so far (L, HN and HC) (8, 43).

**Figure 1.**
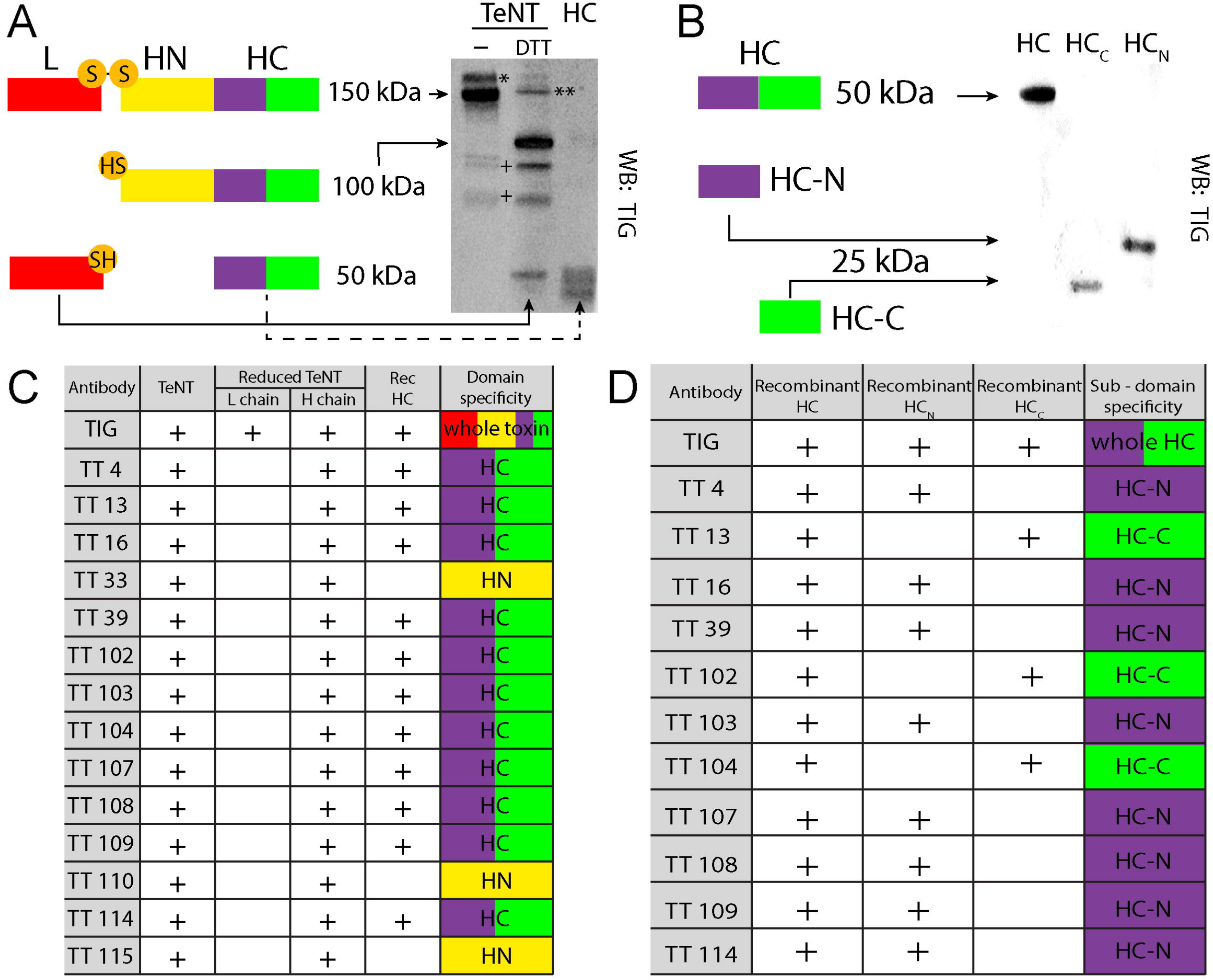
TeNT polypeptide chains and domain-specific recognition by human monoclonal antibodies. A) Schematic structure of TeNT (top left), of the TeNT H chain (middle left, yellow), of the L chain (red) and the HC domain comprising two subdomains, HC-N (violet) and HC-C (green) (lower left); the right part of the panel reports the corresponding western blotting with a hyperimmune human polyclonal antiserum raised by vaccination with tetanus toxoid. Whole TeNT has a molecular weight of 150 kDa corresponding to the L chain + H chain (left line). Upon reduction of the interchain disulfide bond with dithiothreitol (DTT), TeNT displays two bands corresponding to the H chain (HN + HC, 100 kDa) and L chain (L, 50 kDa). The recombinant HC fragment has a molecular weight of approximately 50 kDa. The weak immunoreactivity of the L chain may be due to the low amount of L chain-specific IgGs present in the polyclonal antiserum. * indicates a redox isomer of TeNT (70); ** indicate single-chain TeNT; + indicates degradation toxin fragments. B) Schematic structure of HC (top), HC-N (middle) and HC-C (bottom) and as they appear in western blotting stained with the polyclonal serum. C-D) Summary table of the toxin parts recognized by the TT-humAbs as detected by western blotting of the toxin and of its recombinant HC, HC-N and HC-C.

### Inhibition of TeNT activity by humAbs

HumAbs were tested for their ability to inhibit TeNT entry into the neuronal cytosol of cerebellar granule neurons (CGNs) and cleavage of VAMP-2, a VAMP isoform highly expressed in central nervous system neurons. CGNs were chosen because they are highly susceptible to TeNT and VAMP cleavage is easily testable by immunoblotting and imaging (44). Figure 2A shows that an overnight incubation of CGNs with 50 pM TeNT is sufficient to cleave all VAMP-2 in these neurons. Only four out of the fourteen humAbs tested (TT39, TT104, TT109 and TT110; humAbs:TeNT molar ratio 100:1) were able to prevent TeNT action when preincubated with the toxin, albeit to different extents. Accordingly, we focused on these four humAbs and tested various humAb:TeNT molar ratios to determine their neutralizing potency. As shown in Figure 2B, TT39 and TT109 showed partial neutralization even when added at a 100-fold molar excess of the antibody. In contrast, both TT104 and TT110 prevented TeNT intoxication already at a low molar ratio, TT104 being the most powerful. These results were paralleled by immunofluorescence staining of the VAMP-2 (Figure 2C).

**Figure 2.**
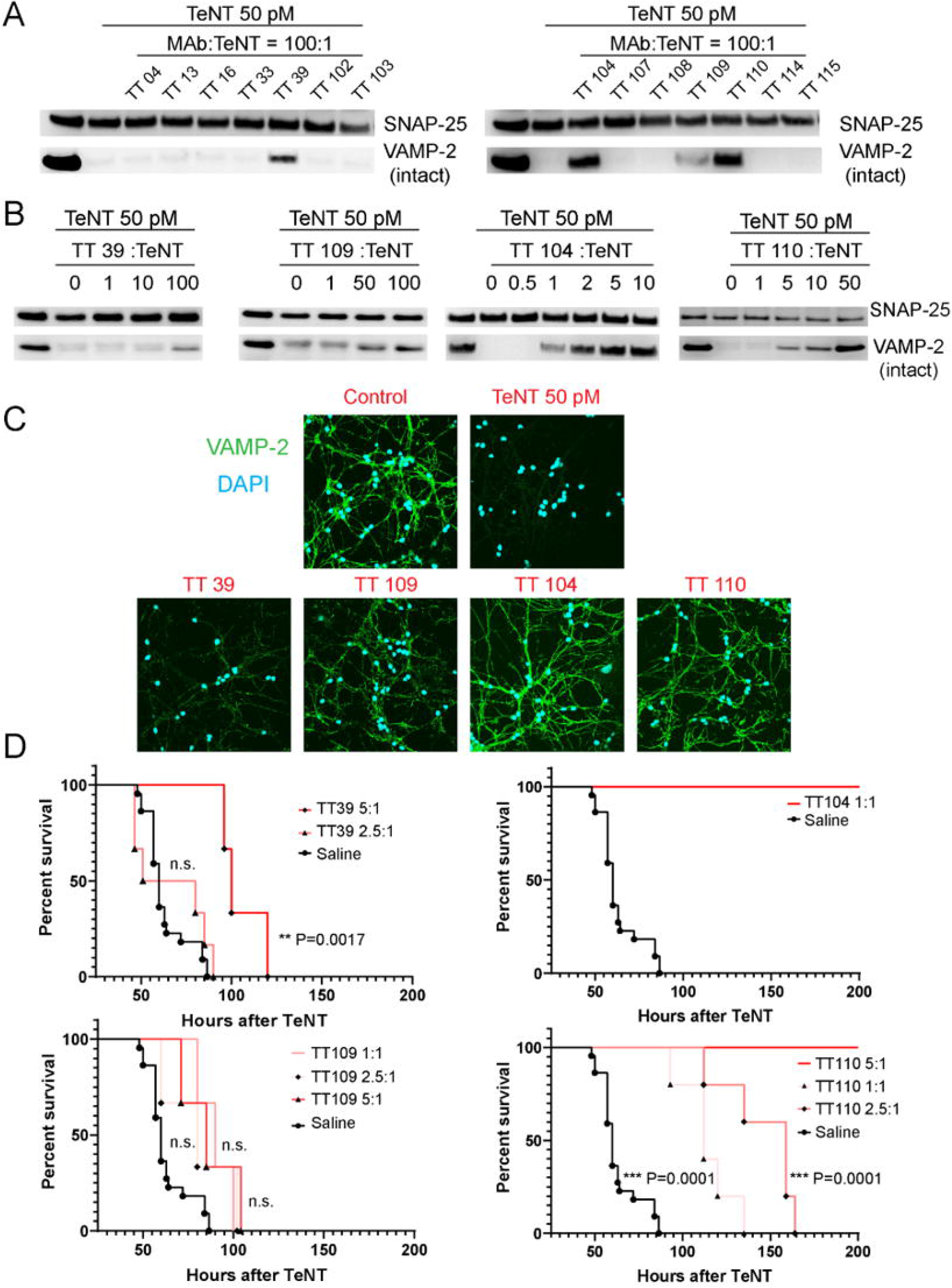
Tetanus toxin neutralization by human mAbs assayed *in vitro* and *in vivo*. A) The initial screening for TeNT *in vitro* neutralization was performed on cultured CGNs that are very sensitive to TeNT. TeNT (50 pM) was diluted in complete culture medium alone or supplemented with a 100X molar excess of the indicated human mAbs (humAbs). The mixture was then added to CGNs for 12 hours. TeNT activity was evaluated by visualizing the disappearance of VAMP-2 with a specific antibody recognizing its intact form. SNAP-25 immunostaining was used as loading control. B) Effect of different TeNT/humAbs ratios for the four humAbs displaying toxin neutralizing activity *in vitro*. C) Immunofluorescence analysis performed with an antibody specific for intact VAMP-2 (green) to assay for the TeNT neutralizing activity of TT39, TT104, TT109 and TT110 preincubated with TeNT (100:1 molar ratio) and added to the primary culture of CGNs. Control CGNs are labelled in green whereas neurons treated with TeNT alone do not display this signal due to the complete cleavage of VAMP by TeNT. CGNs treated with TeNT preincubated with the indicated humAbs display an intermediate signal depending on the neutralization activity of humAbs: only TT104 and TT110 completely prevent toxin entry and cleavage of VAMP-2. D) Mice were injected intraperitoneally with TeNT (4 ng/kg) alone or pre-incubated with the indicated molar ratios humAb:TeNT and the relative mice survivals plotted as a function of time after toxin injection. Statistical significance was calculated with Mantel-Cox test; experiments were performed with at least 6 animals per group.

The selected humAbs were then tested for their ability of neutralizing TeNT *in vivo*. Figure 2D shows that preincubation of TeNT with TT104 and TT110 prevented the development of tetanus in mice in a dose-dependent manner, with TT104 displaying full neutralization at equimolar ratio. On a molar basis, TT110 was less effective, but significantly slowed the appearance of tetanus symptoms at a 2.5:1 molar ratio and completely neutralized TeNT at a 5:1 ratio. In line with the experiments performed in neuronal cultures, TT39 and TT109 did not protect mice from a TeNT challenge. Based on these results, TT104 and TT110 were chosen for further structural and functional analyses.

### Structure of the ternary complex of TeNT with TT104 and TT110 Fab

Recombinant Fab fragments of TT104 and TT110 were produced by introducing a stop codon after CH1 and purified using an anti-CH1 column (Figure S2). The dissociation constants of the toxin-Fab complexes were measured by surface plasmon resonance using immobilized TeNT as a bait and found to be 6.7 × 10^−12^ M and 3.0 × 10^−9^ M for TT104-Fab and of TT110-Fab, respectively (Figure S3). The high affinity of these interactions suggested the possibility that stable binary and ternary complexes could be formed between TeNT and the two Fab. Indeed, immunocomplexes were obtained by incubation of single components either in binary (1:1) or in ternary mixtures (1:1:1) (Figure S4). The ternary [TeNT]-[TT104-Fab]-[TT110-Fab] immunocomplex was purified to homogeneity by gel filtration (total mass 250 kDa). Attempts to crystallize the complex were not successful and therefore we switched to cryo-electron microscopy which provided a structure of the trimeric complex.

As shown in Figure 3, the overall four-domain folding of TeNT is well conserved in the immunocomplex and superimposes with the structures of the whole toxin and isolated domains obtained by X-ray crystallography (8, 45-47). Surprisingly, the trimeric complex [TeNT]-[TT104-Fab]-[TT110-Fab] in solution appears to be quite flexible (Figure S5A), similarly to what observed for TeNT alone.

**Figure 3.**
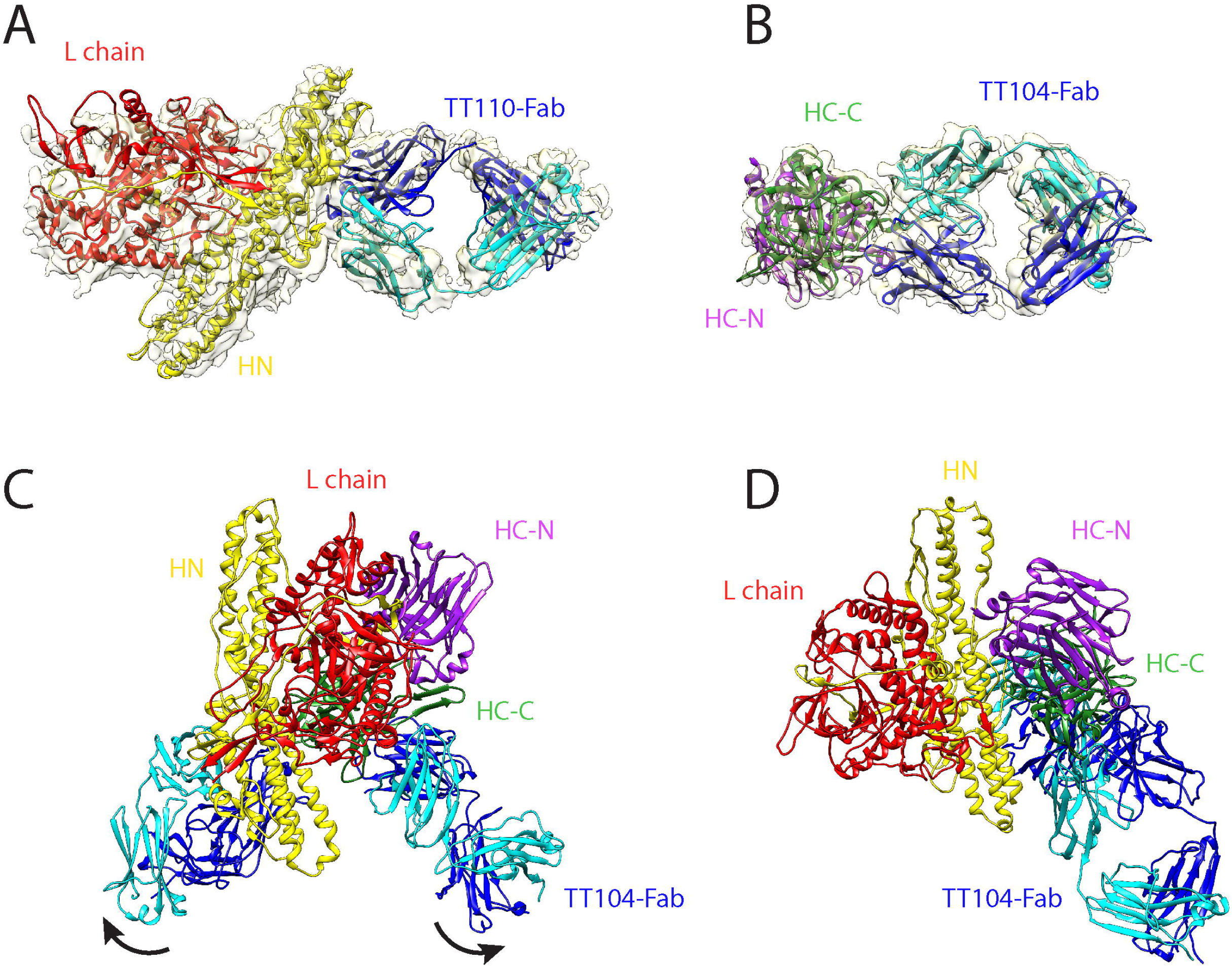
Cryo-EM structure of TeNT-Fabs ternary complex. A) Structure of HN (yellow) and L (red) domains in complex with TT110-Fab (light and dark blue for the variable L and H chain, respectively). B) Structure of the HC-C (green) and HC-N (purple) domains in complex with TT104-Fab (light and dark blue for the variable L and H chain, respectively). C-D) Overall structure of the TeNT-Fabs complex colored as in A and B.

Density map analyses allowed us to localize the two Fabs and TeNT domains, with a clear binding of TT104-Fab to the HC-C domain and TT110-Fab to the HN domain. To overcome the intrinsic flexibility of the complex, the structure was split in two parts, which were treated separately. Masks were generated and used in the multi-body technique to yield two different maps, one relative to HC and TT104-Fab, and the other containing the remaining part of the complex, i.e. TT110-Fab bound to HN linked to the L domain. Interestingly, the overall flexibility is not influenced by Fab binding as it is mainly located around residues 870-875, a loop not resolved in the crystal structure of the toxin and corresponding to the connection between the C-terminal of HN and the N-terminal of HC-N (8).

### Structure of [TeNT-HC]-[TT104-Fab]

The map of the portion of the complex including TT104-Fab and the HC-N and HC-C domains (residues 875-1110 and 1111-1315) has an overall resolution of 3.9 Å and the model could be fitted starting from the available X-ray structure of TeNT (PDB ID 5n0b) and of a Fab modelled on the sequence of TT104-Fab. The area of interaction of the antibody with the HC-C domain is opposite to the HN and L domains and includes portions of polypeptide strands 1140-1145, 1149-1157, 1171-1173, 1202-1204 and 1276-1281 (Figure 3A). From the Fab side, the interaction involves mainly the V_H_ chain whose surface area buried upon the binding is 555 Å^2^, whilst the buried area of chain V_L_ is 396 Å^2^. This difference in the extension of the contact area is reflected in the number of interactions. Residues of chain V_H_ form one salt bridge and seven potential hydrogen-bonds, whilst chain V_L_ contributes to stabilize the protein-protein interaction with additional four hydrogen bonds (Supplementary Table 1, calculation performed with server PISA) (48). The total TeNT buried surface in the complex with TT104-Fab is 792 Å^2^ (Figure 3B). The TT104-Fab binding site is only 12 Å away from the putative nidogen binding site modelled previously (14) and its proximity suggests a possible interference with the binding of nidogen to HC-C due to a steric clash (Figure S6). At the same time, this binding might alter the ability of HC-C to interact with the oligosaccharide portion of PSG, which projects out the neuronal plasma membrane and has to be accommodated into the HC-C to mediate binding (17). These possibilities have been tested experimentally (see below).

### Structure of [TeNT-L-HN]-[TT110-Fab]

The nominal resolution of the L-H_N_-TT110-Fab portion of the immunocomplex (8.3 Å) is lower than that involving TT104-Fab, owing to the preferred orientation assumed by the particles in the grid. However, the overall shape of the TT110-Fab interacting with the H_N_ domain can be clearly distinguished, although the atomic details of the interaction cannot be precisely defined. TT110-Fab binds in the area of the translocation domain HN lying opposite to the L domain (Figure 3).

The overall buried surface of this epitope is estimated to be 700 Å^2^ and 380 Å^2^ for chains V_H_ and V_L_, respectively, and the interaction area involves the helices 597-607 and 614-625, extending to the following strand until residue 631, including segment 655-663. Importantly, this latter segment is part of the “BoNT-switch”, a structural module proposed to be a main driver of the low-pH induced membrane insertion of HN in the botulinum neurotoxins (BoNT), a group of toxins sharing the structural architecture and the mechanism of neuron intoxication of TeNT (49). The BoNT-switch is composed of disordered loops and three short helices (α_A_, α_B_, α_C_) that are also present in TeNT (Figure S7); TT110-Fab binds to a region in TeNT corresponding to the α_B_ of botulinum neurotoxin serotype A1 (BoNT/A1) (Figure S8). At acidic pH, the BoNT-switch rearranges into five β strands (dubbed β1–β5) with α_A_ flipping out from the toxin structure at the center of an elongated hinge formed by β2/β3 hairpins (corresponding to residues I630–Y648 in BoNT/A1 and to V639-Y657 in TeNT). This hinge was shown to insert into the lipid bilayer and to be essential for the subsequent membrane translocation of the L domain of BoNT/A1 (49). At the same time, α_B_ and α_C_ form the β4/β5 hairpin with a hydrophobic surface generated by the structural rearrangement. Considering that TT110-Fab binds to α_B_, it is very likely that this Fab neutralizes TeNT by interfering with the low pH driven insertion of HN into the membrane, thus blocking the translocation of the L domain into the cytosol.

### Model of the overall structure of the immunocomplex

Owing to the intrinsic flexibility of the complex in solution, it was not possible to obtain a complete 3D map of the entire immunocomplex. Nevertheless, thanks to the available crystal structure of TeNT (8, 41), a reliable model of the ternary complex could be built by superimposing the L, HN, HC-N and HC-C domains to the TeNT crystal structure (Figure 3C). It has to be considered that the flexibility of the TeNT-Fabs complex observed in our images is intrinsic to TeNT as it was also observed for the toxin alone using cryo-EM or SAXS measurements (8, 41) and that this model represents one of the possible conformations of the toxin with the two Fabs bound. We detected several eigenvalues (Figure S5B and C) that define the dynamics of the motion of the L-HN domains relative to the HC-N and HC-C domains (Figure 4). All the amplitudes histograms of these eigenvalues were unimodal, suggesting that these motions were continuous. Above them the first three eigenvalue describe 85% of the variability present in the dataset. The movies of the reconstructed body repositioned along these eigenvectors reveals that the two domains of TeNT have a rocking motion (Supplementary movies 1, 2 and 3). Interestingly, the conformation of TeNT in the immunocomplex is closer to the open conformation of the full-length toxin in which the C869-C1093 disulphide bond was reduced (41) than that of the non-reduced TeNT (8).

**Figure 4.**
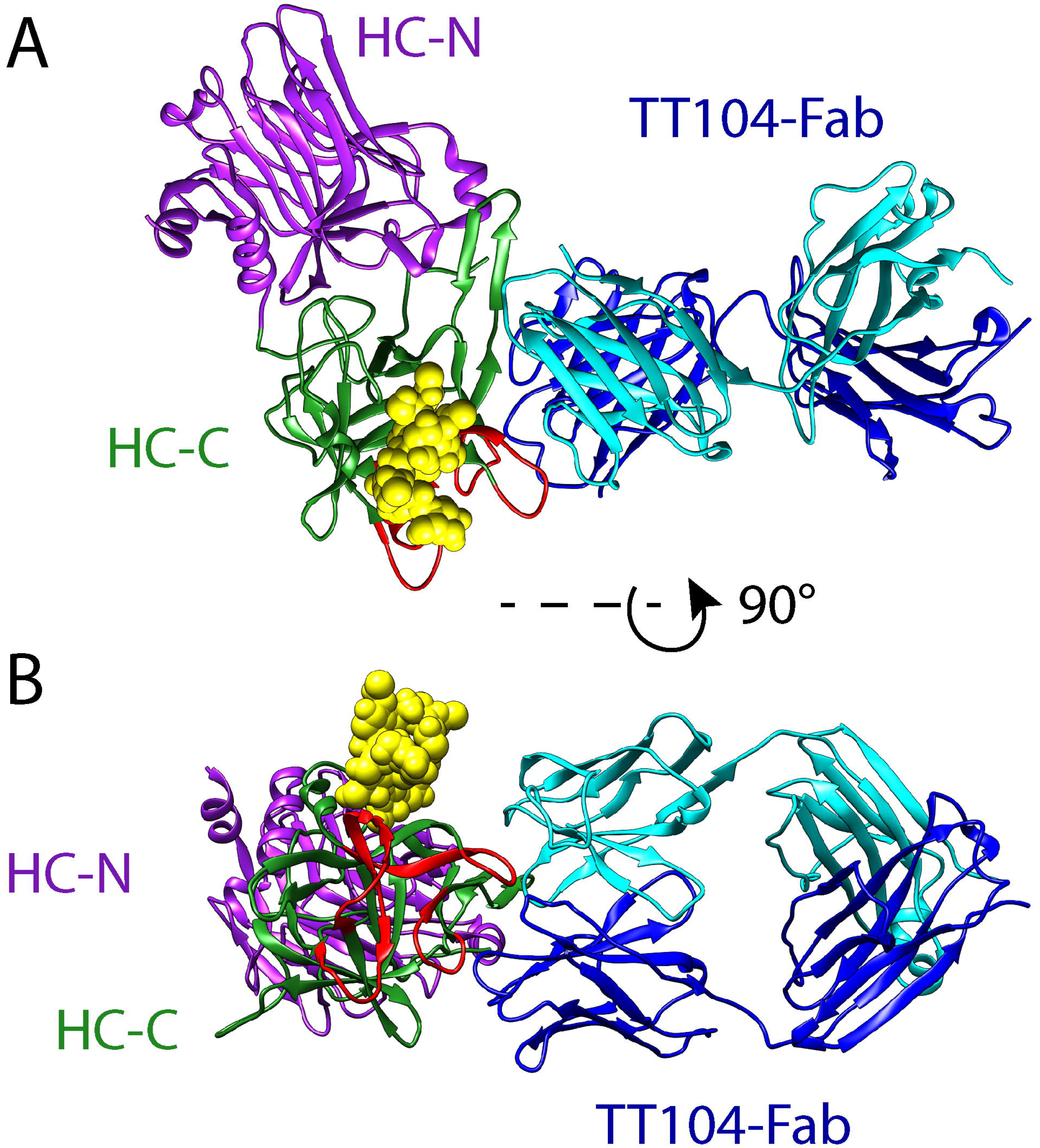
Structure of TT104-Fab bound to TeNT. The HC-N (purple) and HC-C (green) domains of TeNT complexed with TT104-Fab colored as in Figure 3, and the oligosaccharide portion of the GT1b colored in yellow (bound as in pdb:1FV3), and the nidogen binding region highlighted in red.

### TT104-Fab inhibits the biological activity of TeNT by preventing membrane binding

The analysis of the binding between TeNT and TT104-Fab suggests that this Fab should prevent TeNT activity both *in vitro* and *in vivo*. Indeed, TT104-Fab prevents the cleavage of VAMP in cultured CNS neurons and the development of tetanus in mice (Figure 5A and B). These inhibitory effects are caused by the inability of TeNT-TT104-Fab complex to bind the neuronal plasma membrane and the neuromuscular junction (NMJ), as detected using an AlexaFluor555-labelled recombinant HC domain (A555-TeNT-HC, red signal in Figure 5C and D). This inhibition is very specific, as indicated by the lack of effect of TT104-Fab on the binding of a fluorescent HC of BoNT/A1 (CpV-BoNT/A1-HC, green signal in Figure 5C, D), which was pre-mixed with A555-TeNT-HC. The binding domains of the two toxins, TeNT and BoNT/A1, compete among themselves as they both bind PSG (Figure 5C), but TT104-Fab binding to TeNT prevents it from competing with BoNT (Figure 5D).

**Figure 5.**
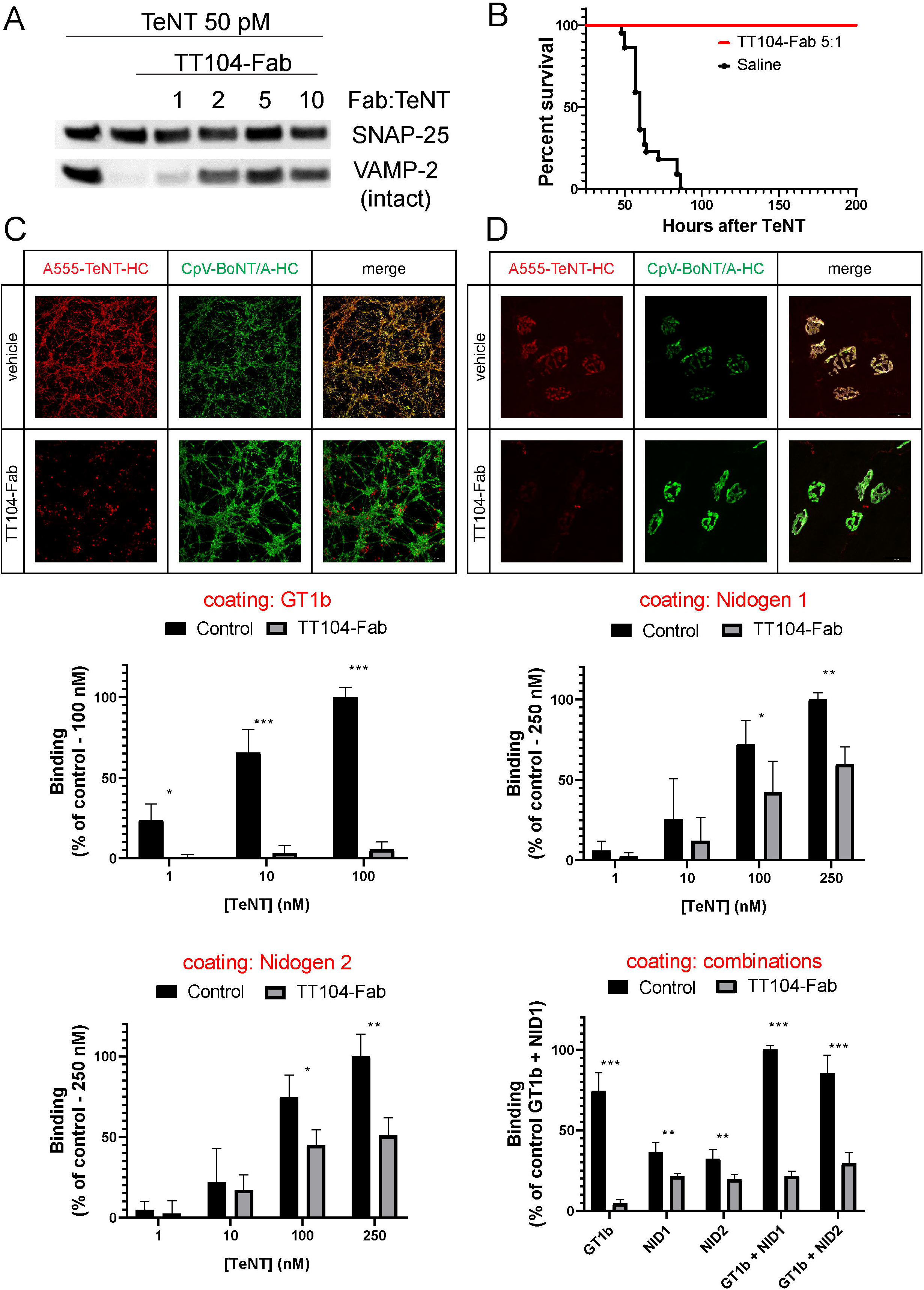
TT104-Fab prevents TeNT toxicity by interfering with its binding to polysialogangliosides and nidogen. A) Western blotting analysis of CGNs treated with 50 pM TeNT preincubated with the indicated TT104-Fab:TeNT molar ratios. After 12 hours CGNs were lysed, electrophoresed and immunoblotted with antibodies specific for VAMP-2 or for SNAP-25 as in Figure 2. B) Survival of mice injected intraperitoneally either with TeNT alone (4 ng/kg, black trace) or premixed with 5:1 molar ratio TT104-Fab:TeNT (red trace). C) Immunofluorescence staining of CGNs treated with a mixture of 50 nM A555-TeNT-HC (red) or 50 nM CpV-BoNT/A-HC (green) preincubated either with culture medium or with a 2:1 molar ratio of TT104-Fab:TeNT-HC for 2 hours and observed with an epifluorescence microscope. Images are representative of experiments performed in triplicates. D) Immunofluorescence staining of the *Levatoris Aureus Longus* muscle injected *in vivo* with A555-TeNT-HC (1 µg) or CpV-BoNT/A-HC (1 µg) preincubated either with vehicle or with a 2:1 molar ratio of TT104-Fab:TeNT and observed after 2 hours with a confocal microscope. Images are representative of a typical experiment performed in triplicates. E) Purified GT_1_b (0.5 μg/well left panel), recombinant nidogen-1/2 (250 ng/well, top right and bottom left panels) or their combination (bottom right panel) were adsorbed by overnight incubation on ELISA plates and the binding of indicated concentrations of either TeNT alone (black columns) or TeNT preincubated with TT110-Fab (gray columns) was tested as described previously (12). Data are reported as percentage of the highest value and averaged from at least three independent experiments. Statistical significance has been calculated using unpaired t tests (*=P<0.05, **=P<0.01; ***P<0.001).

To elucidate the specific mechanism responsible for the TT104-Fab-mediated inhibition of TeNT binding, we assayed the interaction of TeNT with immobilized GT1b, purified nidogen-1 and -2, or their combinations. Figure 5E shows that TeNT binding to GT1b is completely prevented by TT104-Fab, whilst that to nidogens is strongly reduced, with nidogen-1 and nidogen-2 binding affected to a similar extent. This result is in line with the definition of the surface of the conformational epitope recognized by the idiotype of TT104-Fab. Interestingly, when GT1b and nidogens are immobilized together (Figure 5E, right panel), the level of TeNT binding in the presence of TT104-Fab is similar to that reached when only nidogens are present. These findings supports the model of nidogen-TeNT interaction occurring at a distinct site with respect to that of PSG (14), and indicate that TT104-Fab interferes with the binding of both surface receptors. At the same time, these results clearly indicate that membrane binding is the specific step inhibited by TT104 and TT104-Fab, leading to the prediction of a strong anti-tetanus activity *in vivo*.

### TT110-Fab inhibits the low pH induced conformational change of TeNT

Though less powerful than TT104-Fab, also the HN-specific TT110-Fab effectively prevents the cleavage of VAMP by TeNT in cultured neurons (Figure 6A) and protects mice from a TeNT challenge when the toxin and the Fab are pre-incubated together before i.p. injection (Figure 6B). As discussed before, the structure of the immunocomplex suggest that TT110 interferes with the HN-dependent membrane translocation of the L chain driven by acidification. After neurotransmitter release, SV are endocytosed and acidified by the vacuolar ATPase, a process necessary for neurotransmitter re-loading. TeNT, which is internalized inside SV in inhibitory interneurons (21), exploits this physiological process to change structure with exposure of hydrophobic patches on HN that mediate its membrane insertion in such a way as to assist the membrane translocation of the L domain (50). As this process cannot be experimentally accessed from the outside of the cells, we took advantage of a method previously devised to induce the low pH membrane translocation of the L chain directly from the plasma membrane into the cytosol (51-53). As schematized in Figure 6C, after TeNT binding to neurons at 0° C, membrane translocation is induced by replacing the cold medium with acidic medium at 37°C for few minutes. Neurons are then incubated in control medium in the presence of bafilomycin A1, a v-ATPase inhibitor preventing toxin entry through the canonical route, and L domain translocation is assessed by determining VAMP-2 cleavage. Figure 6D shows that upon exposure to pH 5, the L metalloprotease enters the cytosol and efficiently cleaves VAMP-2. In contrast, when TeNT is pre-incubated with TT110-Fab, VAMP-2 is no longer cleaved, consistent with a block of translocation. As discussed above, TT110-Fab could prevent the low pH-induced change of structure of TeNT with ensuing partition of the toxin molecule into the lipid bilayer (50, 54, 55). To test this possibility we performed a fluorometric assay based on the binding of the lipophilic dye ANS to hydrophobic protein patches, thus monitoring the low pH-induced structural change of TeNT (56).

**Figure 6.**
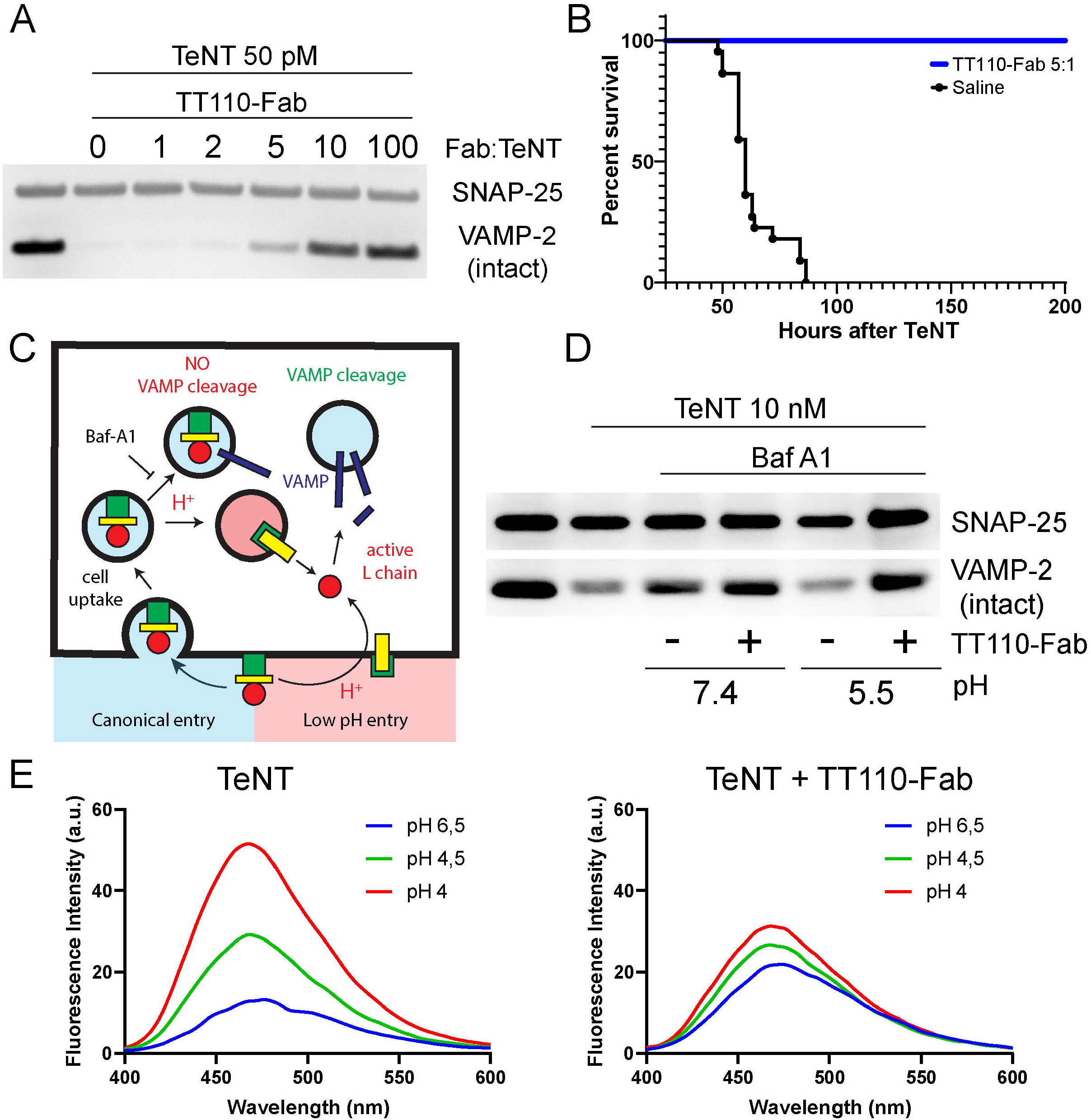
TT110-Fab neutralizes TeNT toxicity by preventing the translocation of the L chain into the nerve terminal cytosol. A) Western blotting of CGNs treated with 50 pM TeNT preincubated with the indicated TT104-Fab:TeNT molar ratio. After 12 hours, CGNs were lysed and immunoblotted for VAMP-2 and SNAP-25 as in Figure 2. B) Survival curve of mice injected intraperitoneally with either TeNT alone (4 ng/kg, black trace) or premixed with 5X molar excess of TT104-Fab (blue trace). C) Scheme illustrating the entry of TeNT L chain into the neuronal cytosol via either: i) the canonical receptor-mediated cell uptake and translocation across the membrane of synaptic vesicles triggered by the acidification of their lumen due to the proton pump activity of the V-ATPase (light blue) or ii) the low-pH translocation induce crossing of the plasma membrane triggered by lowering the culture medium pH in the presence of bafilomycin A1 as previously described (51, 53). D) Western blotting analysis showing the inhibition of TeNT L chain membrane translocation by TT110-Fab. CGNs were incubated at 4 °C for 15 minutes with either TeNT (10 nM) or TeNT preincubated with TT110-Fab. The culture medium was then replaced with a 37 °C buffer for 10 minutes either at pH 7.4 (control condition that does not trigger L chain membrane translocation) or pH 5.0 to trigger the entry of the L chain across the plasma membrane. Samples were then incubated for 12 h with cell culture medium in the presence of bafilomycin A1 (100 nM) and membrane translocation evaluated by assaying the L chain mediate VAMP-2 proteolysis by immunoblotting. SNAP-25 immunoreactivity served as loading control. The image represents the result of a typical experiment. E) ANS fluorescence binding experiment showing the pH-induced conformational change of TeNT blocked by TT110-Fab. TeNT (0.35 µM) (left panel) or TeNT pre-incubated with TT110-Fab (right panel) were incubated at pH 7.0 in the presence of 50 µM ANS and liposomes. The conformational change was triggered by lowering the pH with sequential addition of specific volumes of HCl and evaluated following the ANS fluorescence intensity at 470 nm as previously described (56).

Figure 6E shows that TT110-Fab prevents TeNT from undergoing the low pH-driven conformational change that result in the exposure of the hydrophobic surface. This result supports the hypothesis that TT110 blocks the “BoNT-switch” of HN and prevents the occurrence of the low pH-driven insertion of HN and the following membrane translocation of the L domain. Considering these results and the available literature, we propose that the protonation of Asp618, Asp621, Asp622, Glu626, Glu658, Glu666 of TeNT, is likely to be the key initial event of the low pH conformational transition of HN leading and membrane insertion at acidic pH values. Future mutagenesis experiments targeting these residues will provide conclusive evidence on the role of these acidic sites in the molecular pathogenesis of tetanus.

Collectively, the above experiments demonstrate that TT110-Fab and TT104-Fab act on two different steps of the process of nerve terminal intoxication by TeNT, predicting that they should display additive effects in preventing experimental tetanus *in vivo*.

### Prophylaxis and therapy of experimental tetanus by TT104, TT110 and their Fabs

In the above experiments, the neutralizing activity of TT104 and TT110 was tested by pre-incubating the antibodies with TeNT *in vitro*, a protocol that does not closely match the conditions found in injured patients where IgG anti-TeNT are used as prophylactic agents of TeNT molecules before they are produced by *C. tetani* in necrotic wounds. Therefore, we performed a set of experiments in which mice were pre-treated with a single intraperitoneal injection of TT104 and TT110, alone or in combination, before inoculating 5 MLD_50_ of TeNT ad different delayed time points. Since tetanus usually develops with incubation times ranging between 2 and 15 days after injury (1, 2), animals were pretreated with the TT104 and TT110 alone (400 ng/kg each) or in combination (200 ng/kg + 200 ng/kg) 7 and 15 days before TeNT injection (Figure 7A). As a control, we pretreated a cohort of animals with a human hyperimmune IgG preparation (TIG, 7 IU/kg, corresponding to the standard prophylaxis with 500 IU calculated in a human of 70 kg). Remarkably, TT104 provided full protection of mice from tetanus, even when injected 15 days before TeNT (Figure 7B). TT110 was less effective since it was fully protective from tetanus only when challenge was performed on day 7 (Figure 7C). The combination of 200 ng/kg of TT104 and of 200 ng/kg of TT110 provided a full protection of mice for 15 days, as TIG did (Figure 7D). Once the metalloprotease domain of TeNT has been released inside the cytoplasm of target neurons, the toxin cannot be neutralized any longer by anti-TeNT antibodies. However, worsening of the symptoms can be prevented by neutralization of circulating TeNT, and this is the reason supporting the general practice of injecting TIG into hospitalized patients showing symptomatic tetanus. Thus, we also compared, in a therapeutic setting, TIG and TT104 or TT110 for their capacity to interfere with tetanus at different times after TeNT challenge. We opted to use the Fabs derivatives, which are expected to have a similar activity than humAbs in this type of experimental setting and in consideration of their possible use by intrathecal rather than peripheral administration. To properly compare the Fab and TIG, we estimated the concentration of anti-tetanus specific IgG present in the standard dose of 500 IU (in a putative patient of 70 kg) based on previous quantifications on the serum of hyper-immunized human donors (see methods) (57). As shown in Figure 7E, TeNT inoculation (4 ng/kg) was followed by injection of either TIG or the corresponding amount of TT104-Fab and TT110-Fab in combination after different time periods (1.5, 3, 6 and 12 hours (Figure 7E). Table 1 shows that the two Fab combination completely protected mice, as TIG did, up to 6 hours after injection of TeNT. However, both forms of serotherapy do not block the toxin completely when injected 12 h after TeNT challenge, in agreement with what is known on the kinetics of TeNT internalization into neurons (5). Notably, in this case, mice developed tetanus symptoms with a similar time course (Figure 7F), further indicating that the combination of TT104-Fab and TT110-Fab is as effective as TIG in neutralizing the activity of the toxin present in body fluids.

**Figure 7.**
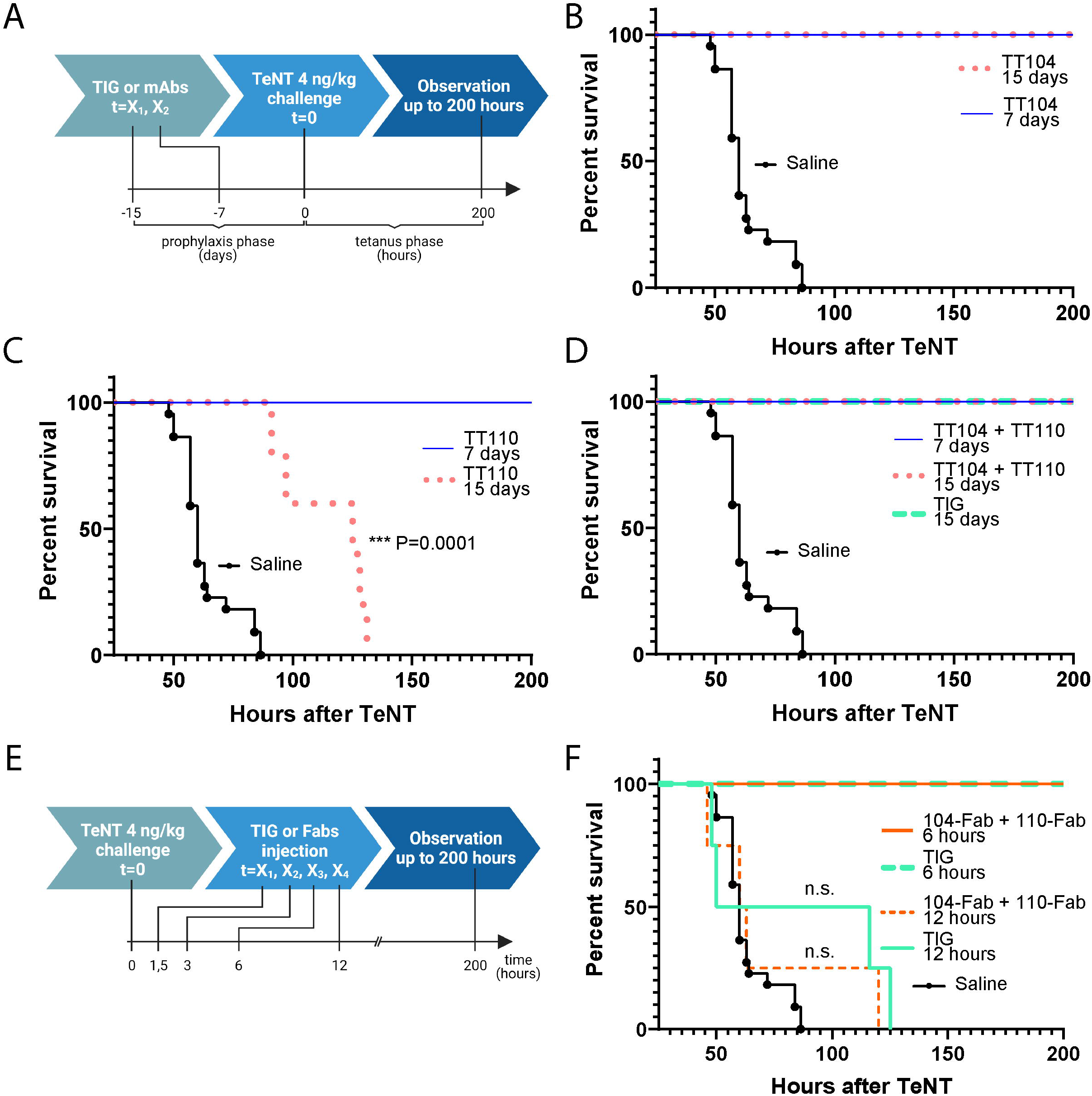
TT104 and TT110 HumAbs allow a long-lasting prophylactic protection against TeNT and their Fabs derivatives prevent tetanus development after toxin challenge. A) Scheme for the time-course to test the prophylactic activity of humAbs. Mice were intraperitoneally pre-injected with either TT104 (400 ng/kg) or TT110 (400 ng/kg) or their combination (200 ng/kg + 200 ng/kg) or with TIG (3.5 IU/kg roughly corresponding to 250 IU/70 kg) for 15 or 7 days. TeNT (4 ng/kg) was then inoculated intra-peritoneally and the animals observed for 200 hours for the development of tetanus symptoms. The prophylactic profiles for TT104 and TT110 injected alone are shown in (B) and (C), respectively. Panel D shows the survival curves of TT104 + TT110 combination compared to TIG, used as a reference. Traces shows the survival curves related to at least 10 animals per group. E) Scheme for the time-course to test the ability of Fabs derivatives to prevent tetanus after TeNT challenge. TeNT (4 ng/kg) was inoculated via intraperitoneal injection. At indicated times, the combination of TT104 + TT110 Fab derivatives (1.2 µg/kg) or TIG (7 IU/kg) were injected intra peritoneum and the animals observed for 200 h for the development of tetanus symptoms. F) Survival plot of mice injected with TeNT and treated with either TT104 + TT110 Fab derivatives (1.2 µg/kg, orange traces) or TIG (7 IU/kg, cyan traces) after 6 or 12 hours. Statistical significance was calculated with Mantel-Cox test; experiments were performed with at least 6 animals per group.

**Table 1:** TeNT neutralization *in vivo* showing the prophylactic and therapeutic activities of HumAbs and Fabs. HumAbs, TIG or Fab were injected at the indicated time points with respect to TeNT inoculation. Survival percentage was calculated based on animals surviving up to 200 hours.

| Prophylactic use |  |  |  |
| --- | --- | --- | --- |
| Agent | Dose | Time before TeNT injection (days) | % survival |
| HumAbs combination | 400 ng/kg | 7 days | 100 % |
|  |  | 15 days | 100 % |
| TIG | 3.5 U/kg (corresponding to 0.6 µg/kg of specific anti-TeNT IgGs and to ~ 2.5 mg/kg of total proteins) | 15 days | 100 % |
| Therapeutic use |  |  |  |
| Agent | Dose | Time after TeNT injection (hours) | % survival |
| Fab combination | 1.2 µg/kg | 1.5 h | 100 % |
|  |  | 3 h | 100 % |
|  |  | 6 h | 100 % |
| TIG | 7 U/kg (corresponding to 1.2 µg/kg of specific anti-TeNT IgGs and to ~ 5 mg/kg of total proteins) | 1.5 h | 100 % |
|  |  | 3 h | 100 % |
|  |  | 6 h | 100 % |

## DISCUSSION

Here we report the structural and functional characterization of two human monoclonal antibodies that neutralize TeNT and prevent tetanus at nearly stoichiometric antibody:toxin ratios, a property maintained by the Fab fragments. The two antibodies bind to two functional domains of TeNT. TT104 binds to the HC-C domain that mediates TeNT binding to neurons, whilst TT110 binds to the HN domain that mediates the translocation of the N terminal metalloprotease domain into the neuronal cytosol.

The structure of the trimeric immunocomplex consisting of TeNT and of the two the Fabs of TT104 and TT110 has been resolved by cryo-EM and the two TeNT epitopes were identified experimentally. This analysis was instrumental to determine the cellular and molecular bases of TT104 and TT110 neutralization activity thus explaining why these antibodies are so potent in preventing tetanus in mice, also after being administered up to 15 days before TeNT. These results are even more relevant considering that the lifetime of human IgGs in mice is expected to be lower than that in humans (58, 59), leading to the prediction that the remarkable duration of protection exerted by TT104 and TT110 will be displayed even longer in humans. It is also noteworthy that TeNT neutralization was effective in the ng/kg range, a dose orders of magnitude lower than the mg/kg range of TIG, offering the possibility of increasing the dose, if necessary. To the best of our knowledge, this is the first report showing such a potent and long-lasting prophylactic effectiveness of anti-TeNT monoclonal antibodies, which fully meets the needs of patients entering hospital emergency rooms with necrotic wound(s) possibly contaminated by *C. tetani* spores. It should also be considered that these antibodies could be produced in a life extended form in order to improve their half-life and bio-distribution (60). TT104 and TT110 are expected to be very effective also in preventing maternal and/or neonatal tetanus in the case of delivery from a mother not immunized with the tetanus toxoid, a frequent condition in some parts of the world (30).

The currently used serotherapy with human anti-tetanus immunoglobulins (TIG), prepared from hyperimmune donors, is affected by a number of drawbacks such as: i) variation of neutralizing power from lot to lot; ii) possible risk of contamination due to the source of antibodies, iii) need of injecting relatively large amounts of proteins, particularly in a therapeutic setting (30) and, iv) the risk of anaphylactic reactions or serum sickness in the case of horse antisera, v) in addition, patients with a IgA deficit could mount an immune response to the small amount of IgA present in TIG. All these problems could be overcome by the injection of small amounts of well characterized monoclonal antibodies such as those identified in this study or even a of single one, as exemplified by TT104. Our findings also pave the way to the use of monoclonal antibodies and their Fab fragments for administration via the intrathecal route. This procedure was found to provide better results than peripheral injections when the toxin has already undergone retroaxonal transport and release into the spinal cord but is strictly limited by the amount of protein that can be safely injected intrathecally (32, 33).

Recently, two groups reported on the isolation of anti-TeNT humAbs from human B memory cells. A first study showed that tetanus neutralization in mice could be achieved only by preincubation of the toxin with a combination of 3 antibodies used at a fifteen-fold molar excess (42). In another study TeNT neutralization was achieved only at a 3,000-fold antibody molar excess (61). In these cases, as in our study, the main neutralizing effect was provided by antibodies specific for the HC binding domain of TeNT. This notion agrees with the numerous reports about the activity of monoclonal antibodies specific for the HC portion of TeNT isolated from mice, or prepared after humanization procedures, or by scFv antibody phage display library (61-63). Together with our findings with TT104, these data fit with the rather general rule that neutralizing antibodies interfere with pathogen binding to its receptor. However, it is interesting to note that the strong neutralizing activity of TT110 indicates that antibodies can also effectively neutralize TeNT by blocking its membrane translocation.

Nine of the humAbs identified in the present study recognize epitopes localized on the HC-N domain, three bind the HC-C domain and three bind the HN domain. These results support recent evidence that HC is the most immunogenic part of TeNT (42, 61), and extend this finding by suggesting that the most immunogenic domain of HC may be HC-N rather than HC-C. In addition, the fact that HC-N specific humAbs have negligible neutralization activity suggests that the role of HC-N domain is likely to be only structural. This interpretation is in line with the result of Masuyer et al., who showed a direct interaction at low pH of the L and HC-C domains, but not with the HC-N (8).

In addition to the biomedical properties of the two anti-TeNT humAbs, the structure of the immunocomplex formed by the toxin with the two Fab derivatives brought novel information on two key steps of the mechanism of nerve terminal intoxication by TeNT. The first insight is about the sequential events leading to a productive binding of TeNT to the presynaptic membrane at the NMJ. TeNT binds nerve terminals via the interaction with two receptors, the polysialoganglioside and a glycoprotein molecule, identified as the basal membrane protein nidogen 1/2. However, the sequence of binding events is not known. Nidogens are particularly enriched in the NMJ basal lamina which enwraps perisynaptic Schwann cells and the nerve terminal secluding them from the muscle fiber and is permeable to molecule as large as IgG and TeNT. Nidogens are in a strategic position to capture TeNT molecules entering the NMJ, because they are exposed to the perineural extracellular fluids, yet they are engaged in multiple protein-protein interactions with other basal lamina components, which may limit their availability for TeNT fixation. PSG are glycolipids located on the outer leaflet of the presynaptic membrane, which are endowed with long and flexible headgroups projecting above the plasma membrane, as confirmed by their established role as antigens for autoantibodies in Guillain-Barre syndrome (64-68). Our data show that TT104 fully disrupts TeNT interaction with PSG but only partially with nidogens, yet this is sufficient to completely abrogate TeNT binding and internalization at nerve terminals and prevent tetanus. This strongly suggests that the interaction with PSG is a prerequisite for TeNT toxicity and it is the first step of TeNT binding. Nidogens might then drive the TeNT-PSG complex towards the specific endocytic organelle leading to the formation of signaling endosomes (69).

Experiments with TT110 provided new insights on the mechanism by which the HN initiates its membrane insertion to translocate the L domain in the cytosol. TT110-Fab was found to prevent the low pH-driven conformational change of HN by binding to an epitope at the center of an important structural-functional module, the BoNT-switch, previously identified in botulinum neurotoxins serotype A1, which was suggested to act as the pH sensor triggering the initial event of HN membrane insertion (49). The present work indicates that such a structural switch is present also in TeNT. In addition, the TeNT epitope recognized by TT110 identifies a group of carboxylate residues that are candidates for future studies aimed at determining their role in the TeNT low pH triggered switch for membrane insertion.

The results described here qualify the humAbs TT104, TT110 and their Fabs derivatives as novel therapeutics highly superior to the presently used anti-tetanus immunoglobulins, potentially ready, after appropriate formulation, to replace the immunoglobulins purified from human or equine blood in clinical practice. These anti-TeNT humAbs and their Fab derivatives open new avenues to the treatment of tetanus patients because large doses could be injected intrathecally, thus maximizing the ability of these novel therapeutics to neutralize TeNT in the very area of its action, i.e. the spinal cord.

## ACKNOWLEDGEMENTS

We are grateful to the European Synchrotron Radiation Facility in Grenoble (France) for provision of beam time on CM01. We thank Gordon A. Leonard for the support in data collection. We thank Prof. Ornella Rossetto for critical reading of the manuscript.

We gratefully acknowledge the financial support of the University of Padova (M.P. and C.M.), of the project RIPANE (C.M.), the Wellcome Trust (107116/Z/15/Z) [G.S.] and the UK Dementia Research Institute (UKDRI-1005) [G.S.].

## AUTHOR CONTRIBUTIONS

Conceptualization, C.M., A.L. and G.Z.; Investigation M.P., A.G., O.L., F.V., M.T. S.B, D.C., C. SF., E.K., L.O.; Data Curation, A.G. and G.Z., Resources, G.S., S.B., D.C., C. SF., E.K., L.O.; Supervision, C.M., A.L. and G.Z. Writing Original Draft, C.M., A.L. G.Z., G.S. and M.P.; Writing, Review and Editing, all authors.

## DECLARATION OF INTERESTS

A.L., L.P., C.S. and D.C. are employees of VIR Biotechnology and may hold shares in Vir Biotechnology.

## Material and methods

### Mice and *in vivo* procedures

Swiss-Webster adult female CD1 mice (Charles River Laboratories) were housed under controlled light/dark conditions and maintained on a regular diet of food and water provided *ad libitum*. All experiments were performed in accordance with the European Community Council Directive n° 2010/63/UE and approved by the Italian Ministry of Health. For experimental tetanus assays, animals were randomly assigned to test and control groups. TeNT, humAbs, Fabs or commercial tetanus immunoglobulin (TIG) were administered by intraperitoneal (IP) injection at doses and modalities described in the method details.

### Cell cultures of cerebellar granule neurons

Primary cultures of rat cerebellar granule neurons (CGNs) were prepared from 4-to 6-day-old rats as previously described (44). Briefly, rat cerebella were isolated, mechanically disrupted and trypsinized in the presence of DNase I. Cells were then collected and plated into 24-well plates pre-coated with poly-L-lysine (50 μg/ml) at a cell density of 4×10^5^ cells per well or onto 13 mm round glasses at a cell density of 3×10^5^ cells per well. Cultures were maintained at 37°C, 5% CO2, 95% humidity in BME (Basal Medium Eagle) supplemented with 10% fetal bovine serum, 25 mM KCl, 2 mM glutamine and 50 μg/ml gentamicin (here after indicated as complete culture medium). To arrest growth of non-neuronal cells, cytosine arabinoside (10 μM) was added to the complete culture medium 18–24 h after plating. Neurons were used after 6 days *in vitro* and to a maximum of 8 days *in vitro*.

### Reagents and proteins

All chemicals used were from Sigma Aldrich. TeNT was previously isolated from culture filtrates of *C. tetani* strain Y-IV-3 (WS 15), frozen in liquid nitrogen and stored at -80°C in 10 mM HEPES-NaOH, 50 mM sodium chloride, pH 7.2.

TeNT HC (residues 875-1,315) was N-terminally fused with glutathione *S*-transferase (GST) and expressed in *E. coli* as previously described (71). Where indicated, TeNT HC was labeled with AlexaFluor555-maleimide following manufacturer’s instructions and dialyzed against 10 mM HEPES-NaOH, 150 mM NaCl, pH 7.4. TeNT HC-N (residues 856-1,110) and HC-C (residues 1,111-1,315) were expressed in *E. coli* as GST fusion proteins and purified as previously described (72). HC BoNT/A1 (residues 876-1296) was N-terminally fused with CpV (Circularly permutated Venus), cloned into a pET28a His-tag vector (Novagen) and expressed in *E. coli*. Purification was performed with a prepacked HisTrap Ni column (GE Healthcare) as previously described (73). Protein concentration was determined by absorption spectroscopy and purity by SDS-PAGE electrophoresis on 4-12% NuPAGE gels in MES buffer (Life Technologies).

### Human monoclonal antibodies and Fab fragments

Memory B cells were isolated from peripheral blood mononuclear cells (PBMCs) through magnetic cell sorting with 0.5 μg/ml anti-CD19-PECy7 antibodies (BD, 341113) and mouse anti-PE microbeads (Miltenyi Biotec, 130-048-081) followed by FACS sorting using 3.75 μg/ml Alexa Fluor 647–conjugated goat anti-human IgG (Jackson ImmunoResearch, 109-606-170), 5 μg/ml Alexa Fluor 647–conjugated goat anti-human IgM (Invitrogen, A21215) and PE–labeled anti-human IgD (used at a 1:40 dilution; BD Biosciences, 555779). As previously described (35), sorted B cells were immortalized with Epstein–Barr virus (EBV) and plated in single-cell cultures in the presence of CpG-DNA (2.5 μg/ml) and irradiated PBMC-feeder cells. Two weeks post-immortalization, the culture supernatants were tested (at a 2:5 dilution) for binding to TT by ELISA. Briefly, ELISA plates were coated with 1 μg/ml of recombinant TT. Plates were blocked with 1% BSA and incubated with titrated antibodies, followed by 1/500 alkaline phosphatase (AP)-conjugated goat anti– human IgG (Southern Biotech, 2040-04). Plates were then washed, substrate (para-nitrophenyl phosphate (p-NPP), Sigma) was added and plates were read at 405 nm.

Recombinant Fab fragments were produced in HEK cells by introducing a stop codon at the end of CH1 and purified by affinity chromatography on AKTA Xpress Mab System (Cytiva) with UNICORN 5.11 software version (Build 407) using CaptureSelect CH1-XL MiniChrom columns (ThermoFisher Scientific), buffer exchanged to PBS using a HiPrep 26/10 desalting columns (Cytiva). Purified FABs were concentrated by Amicon Ultra filter units (Millipore), sterilized through a 0.22 mm filter and stored at -80°C after rapid freezing in liquid N2.

### TeNT-Fabs complexes formation and sample purification

TeNT and the two Fabs were incubated at 4°C overnight under stirring. The complex formed was purified from unbound Fabs by gel filtration (Superdex 200 10/300, GE) and analyzed by a Native-PAGE analysis in buffer 10 mM Tris, 150 mM NaCl, pH 7.4 and visualized in 4-16% native-PAGE gel (ThermoFischer Scientific). The fractions containing the ternary complex were collected and concentrated until a final volume of 500 µl, and subsequently loaded on a gel filtration column. The first peak corresponds to complex TeNT-Fabs (250 kDa) and the second peak at lower molecular weight corresponds to the unbound Fabs (104 and 110). The ternary complex was purified by gel filtration. Native-PAGE gel confirmed that in the peak are present both TeNT, TT104-Fab and TT110-Fab and that the ternary immunocomplex eluted as a homogeneous physical species. At variance, the second peak contained the unbound Fabs, which elute at the same elution volume owing to the same molecular size (data not shown). The peak corresponding to [TeNT]-[TT104-Fab]-[TT110-Fab] was isolated and the complex was concentrated to 1 mg/ml.

### Surface Plasma Resonance (SPR) analysis

For the SPR analysis, a BIAcore™ T100 system (GE Healthcare) was used. TeNT was covalently coupled to a CM5 (series S) sensor chip (carboxymethylated dextran surface) by amine-coupling chemistry to a final density of 1000 resonance units, as previously described (74). A 10 mM acetate pH 5.0 buffer was used for the immobilization. A flow cell with no immobilized protein was used as a control. Binding analysis was carried out in a running buffer consisting of 10 mM Hepes-NaOH, pH 7.6, 150 mM NaCl, 2 mM MgCl_2_, using a flow rate of 30 μl/min. Each sensogram (time course of the surface plasmon resonance signal) was corrected for the response obtained in the control flow cell and normalized to baseline. After each injection, the surface was regenerated by a double injection of 1M NaCl for 1 min to restore the base line to the initial resonance unit value. For kinetics experiments, a Biacore method program was used that included a series of three start-up injections (running buffer), zero control (running buffer), and five different dilutions of the TT104-Fab (500, 200, 50, 10, 2 nM) and four for TT110-Fab (500, 200,100, 50 nM). Serial dilutions were performed in running buffer. High performance injection parameters were used. The contact time was 180 s followed by a 200 s dissociation phase. The kinetic data were analyzed using the 2.0.3 BIAevaluation software (GE Healthcare). The curves (both association and dissociation phases) were fitted with the classical Langmuir 1:1 model; the quality of the fits was assessed by visual inspection of the fitted data and their residuals, and by chi-squared values. Two independent experiments in triplicates were performed.

### Cryo-EM data collection and processing

A first screening of the sample and its analysis with Cryo-EF (75) suggested a strong particle’s orientation (data not shown). In order to overcome this issue, we performed two data acquisitions: a planar one and one with a tilt angle of 30°. For the planar acquisition, 3 ml of the sample at 0.5 mg/ml concentration, were applied to glow-discharged C-flat 2/1 -3 Au holey grid and vitrified in a Mark IV Vitrobot (FEI), whereas, for the tilted acquisition, 3 μl of the sample at 0.35 mg/ml concentration were applied to glow-discharged Aultrafoil 1.2/1.3 grid. Both the grids were imaged in a Titan Krios microscope (Thermo Fisher Scientific) at 300 keV with a K2 direct electron camera at 0.827 Å per pixel. The planar dataset is composed by 402 movies (40 frames each and a dose of 1.2 e^-^/Å^2^) and was combined with the tilted dataset containing 2970 movies (50 frames each with 0.68 e^-^/Å^2^) after beam-induced motion correction with Motioncor2 (76) and Contrast transfer function (CTF) estimated with Gctf (77). The selected micrographs were picked and analyzed in RELION-3 (78). An initial model generated with EMAN2 (79) was used for the 3D classification and refinement. An analysis of the 2D classes and of the obtained 3D model suggests the presence of some mobility of the domains of the toxin. To overcome this problem, a multibody refinement procedure was performed (80). This refinement gives twelve motion eigenvalues with the first three that explain the 85% of the variance in the data. These eigenvalues allow one to align and subtract the signal of the two domains of the protein and to perform a local reconstruction of the two main flexible regions (81, 82) (see workflow in Figure S5). This protocol yielded a 3.9 Å resolution map that matches to the density of the HC domain and the TT104-Fab formed by 98170 particles, and a 8 Å low-resolution map of the LC-HN domains bound to TT110-Fab.

### Model building and refinement

The two maps were visualized and treated separately because of their different resolutions. The crystal structure of TeNT (8) (PDB ID 5n0b) was divided in two files, one including only the HC domain, the domain HN and domain L. The homology models of the structure of the two Fabs were generated via SwissModel (83) using the PDB ID 5dk3 e 1hzh as a template. All models were fitted as rigid bodies using Chimera (84). The HC and TT104-Fab models were fitted in the appropriate map. For the Fab, the two positions rotated by 180° around the molecular pseudo-symmetry axis were tested, and the positions that gave significantly better interactions with the toxin were used. The model was improved by iterative cycles of Phenix real space refinement (85) and visualized with Coot (86). Model refinement was carried out using geometry restraints, including secondary structure, rotamer and Ramachandran plot restraints. Finally, the map was sharpened according with the PDB model’s B-factor using Phenix. The final statistics on the refined models are reported in Supplementary Table 2.

The HN and L chain domains and TT110-Fab were fitted in the second map; owing to the remaining preferential orientation symptom, the quality of the map allows only to perform a rigid body fitting of the already existing structures (8).

### HumAbs subdomain specificity

Whole TeNT (0.5 μg), reduced TeNT (0.5 μg in 15 mM of dithiothreitol), HC-T (1 μg) or the individual subdomains (HC-C and HC-N) were separated by SDS-PAGE electrophoresis on 4-12% NuPAGE gels in MES buffer and transferred onto Protran nitrocellulose membranes. Saturation was carried out in 5% BSA in PBS containing 0.5% of Tween (PBS-T) for 1 hour. HumAbs were diluted in saturation solution at a final concentration of 1 ug/ml and used as primary antibodies. After overnight incubation at 4 °C, membranes were rinsed in PBS-T and incubated with anti-human secondary antibodies conjugated to HRP for 1 hour at room temperature. Membranes were extensively washed and revealed with an Uvitec gel documentation system (Cleaver Scientific) using Luminata Crescendo (Millipore) as substrate.

### TeNT neutralization assay on primary neuronal cultures

CGNs at 6–8 days in vitro (DIV) were treated with TeNT (50 pM) alone or pre-incubated for 1 hour at room temperature at the indicated molar ratios of TeNT:humAbs or TeNT:Fabs in complete BME. After 12 hours, CGNs were lysed with Laemmli sample buffer containing protease inhibitors (Roche) or fixed for 10 min with 4% (w/v) paraformaldehyde in PBS. Neuronal lysates were loaded onto NuPage 12% Bis-Tris gels (Life technologies), separated by electrophoresis in MES buffer and transferred onto Protran nitrocellulose membranes for immunoblotting. Neutralization was determined by monitoring the cleavage of VAMP-2 with a primary antibody specific for the intact form of the protein (Synaptic System 104 211) and an antibody specific for SNAP-25 (clone SMI81 Biolegend) as loading control. Protein amount was revealed either with an Odyssey imaging system (LI-COR Bioscience) or with an Uvitec gel doc system (Uvitec Cambridge) using appropriate antibodies.

Fixed CGNs were analyzed by immunofluorescence. After fixation, CGNs were quenched (50 mM NH_4_Cl in PBS) for 20 min, permeabilized with 0.3% Triton-X100 for 3 min, saturated with PBS 2% BSA for 1 hour at RT and stained with primary antibodies specific for whole VAMP-2 (Synaptic System 104 211). After extensive washing with PBS 0.5% BSA, primary antibodies were detected with appropriate secondary antibodies conjugated to Alexa fluorophores and nuclei were stained with DAPI. Coverslips were mounted using Fluorescent Mounting Medium (Dako) and examined with a Leica SP5 confocal microscope (Leica Micro-systems, Wetzlar, Germany).

### Internalization assay of fluorescent TeNT and BoNT/A HC derivatives in cultured neurons and in mice

BoNT/A-HC-CpV (100 nM) and TeNT-HC-555 (20 nM) were mixed for 1 hour at 37°C in complete BME with or without TT104-Fab (30 nM), and then added onto CGNs at 6-8 DIV seeded on glass coverslips. At indicated time points, cells were gently washed with fresh complete BME medium and incubation stopped with 4% (w/v) PFA. Coverslips were extensively washed and mounted using an appropriate Mounting Medium (Dako). Fluorescence was examined with a Leica SP5 confocal microscope.

For imaging studies in mice, TeNT-HC-555 (1 µg) was mixed with either BoNT-HC-CpV (1 µg) or an equivolume of PBS in saline solution (0.9 % NaCl, 0.2 % gelatin). These solutions were split and incubated for 1 hour at 37 °C with TT104-Fab (1:5 final molar ratio with TeNT-HC-555) or the corresponding volume of saline solution. The mixture was locally injected (final volume of 20 µl) below the skin at the level of the *Levator Auris Longus* (LAL) muscle of anesthetized CD1 mice. After 2 hours, animals were euthanized, the LAL was dissected as previously described and fixed with 4% PFA for 15 minutes (87). The muscles were then directly mounted in Mounting Medium and the binding and internalization of HC probes was evaluated with a Leica SP5 confocal microscope.

### Binding assay to polysialoganglioside GT1b and nidogens

96-well-plates polystyrene plates (Sarstedt, Nümbrecht, Germany) were coated with either 1 µg of GT1b (Santa Cruz Biotechnology, Dallas, TX, USA) dissolved in methanol, or 250 ng of nidogen 1/2 (Bio-Techne, Minneapolis, MN, USA) diluted in PBS, or their combination thereof and let to dry overnight at room temperature. Wells were washed with PBS-T, blocked with 1% BSA in PBS for 1 h at room temperature, and the indicated concentrations of TeNT diluted in PBS, with or without pre-incubation with TT104-Fab (1 hour at RT) were added for 2 hours at RT. Wells were then extensively washed with PBS-T and incubated with a rabbit TeNT antiserum (Istituto Superiore di Sanità, Rome, Italy) diluted in 1% BSA in PBS for 1 hour. After washing with PBST, wells were incubated with appropriate secondary antibody conjugated with HRP. Wells were extensively washed and 100 µL of 2,2′-azino-bis(3-ethylbenzothiazoline-6-sulfonic acid). Absorbance was read at 450 nm with a microplate reader (Tecan).

### Low pH membrane translocation of TeNT in CGNs

CGNs were plated into 24-well plates at 4×10^5^ density. At 6–8 DIV, neurons were incubated in ice-cooled complete BME with either 10 nM TeNT or 10 nM TeNT preincubated with 20 nM TT110-Fab and kept at 4 °C for 15 minutes. Thereafter, pre-warmed medium A (123 mM NaCl, 6 mM KCl, 0.8 mM MgCl2, 1.5 mM CaCl2, 5 mM NaPi, 5 mM citric acid, 5.6 mM glucose, 10 mM NH_4_Cl), adjusted either at pH 7.4 or 5.5 with 1 M TRIS-OH, was added and left for 10 minutes at 37°C. Cells where then washed twice with MEM and incubated in complete BME containing 100 nM Bafilomycin A1 for 12 hours. Translocation of TeNT L chain into the cytosol was evaluated by western blotting following its VAMP-2 specific proteolytic activity as described above.

### Fluorescence assay of ANS binding to clostridial neurotoxins as a function of pH

This assay was performed as previously described (56). Briefly, DML (1,2-dimyristoyl-*sn*-glycero-3-phosphocholine monohydrate) and DMPA (1,2-dimyristoyl-*sn*-glycero-3-phosphate mono sodium salt) in the ratio 4:1 were dissolved in chloroform and methanol (3:1 volume/volume ratio) and dried in a rotavapor system. The lipid film was then rehydrated in double-distilled water and ultrasonically dispersed for 30 minutes at 25 °C in an Ultrasonic bath sonicator (Falc, Italy). The liposomal suspension was homogenized to 100 nm unilamellar vesicles with an extruder and appropriate membranes (Avanti Polar Lipids), yielding a clear suspension.

TeNT alone or TeNT preincubated with tenfold TT110-Fab was diluted to the final concentration of

0.8 μM in 100 mM TRIS-citrate buffer, 100 mM NaCl, pH 7.0 in the presence of liposomes (final concentration of 0.4 mM) and 50 μM ANS. The indicated pH values were reached by adding specific volumes of HCl (6N) directly in the cuvette. Changes in ANS fluorescence were determined at 25°C in a Perkin Elmer LS50B Spectrometer at a scan speed of 800 nm min^−1^ with excitation slit of 15 nm and emission slit of 10 nm following emission between 400 and 650 nm upon excitation at 380 nm. Samples were kept at 25 °C for 10 min before measurements and fluorescence intensities were estimated by averaging five readings. Blanks were mathematically subtracted from spectra.

### Mouse bioassay and antibody neutralization assay

Swiss-Webster adult CD1 mice were housed under controlled light/dark conditions, and food and water were provided ad libitum. All experiments were performed in accordance with the European Community Council Directive n° 2010/63/UE and approved by the Italian Ministry of Health.

For the assay of the neutralization ability of humAbs, TeNT was diluted at a concentration of 8 pg/μl in “bioassay solution” (0.9% NaCl, 0.2% gelatin), split into aliquots and supplemented either with an equivolume of bioassay solution (positive control) or an equivolume of bioassay solution supplemented with the indicated amounts of humAbs or Fabs. These solutions were kept at room temperature under a gentle agitation for 1 h. Female mice (24–26 grams) were randomly injected intra peritoneum with 1 μl per gram of body weight of either toxin alone or toxin-humAbs/Fabs solutions. The final TeNT dose was 4 ng/kg, roughly corresponding to a fivefold lethal dose of our toxin preparation.

For the prophylactic activity, TIG (Igantet, Istituto Grifols Poligono Levante S.A., Italy) or humAbs (TT104 and TT110 alone or their combination) were diluted in bioassay solution and intraperitoneally injected with the indicated dose. After 7 or 15 days, TeNT (4 ng/kg) was injected intraperitoneally and mice were monitored every 6 hours for the first 48 hours and then every 4 hours up to 200 hours, when the experiment was terminated. A human endpoint was set as to when mice showed moderate tetanus symptoms (hunched back and paralysis of rear limbs or disappearance of the righting reflex for 30 s), after which the animal was euthanized by cervical dislocation and considered positive for tetanus. TIG was used at a dose of 7 IU/kg, roughly corresponding to the canonic prophylactic injection of 500 IU in a human of 70 kg.

To test the post-exposure effect of Fabs, TeNT (4 ng/kg) was injected intraperitoneally. At indicated time points, TT104-Fab in combination with TT110-Fab (1.2 µg/kg) or TIG (7 IU/kg) were injected intra peritoneum and mice monitored up to 200 hours. TIG dose was chosen to correspond to the suggested treatment of 500 IU in human of 70 kg. Fab doses were calculated accordingly, estimating the concentration of TeNT-specific IgGs in the serum of hyper-immunized subject equal to 40 µg/ml as averaged from values reported in (57)

Data were plotted as Kaplan-Meier survival curves and statistic measured with the Log-rank test. Each curve is representative of at least 6 animals. At each experiment at least 3 mice were treated with TeNT alone, and the curve for survival of TeNT alone derives from all the data plotted together.

### Data deposition

Atomic coordinates and density maps have been deposited for immediate release as PDB ID 7OH0 and EMD-12890 for the HC-TT104-Fab region and as PDB ID 7OH1 and EMD-12891 for the L-HN-TT110-Fab region

