## Supplementary figures for "Extremely potent Human Monoclonal Antibodies for the Prophylaxis and Therapy of Tetanus"

**Figure S1**: Representative western blotting of TeNT polypeptide chains, domains and sub-domains recognition by the repertoire of human monoclonal antibodies produced by immortalized clonal B memory cells

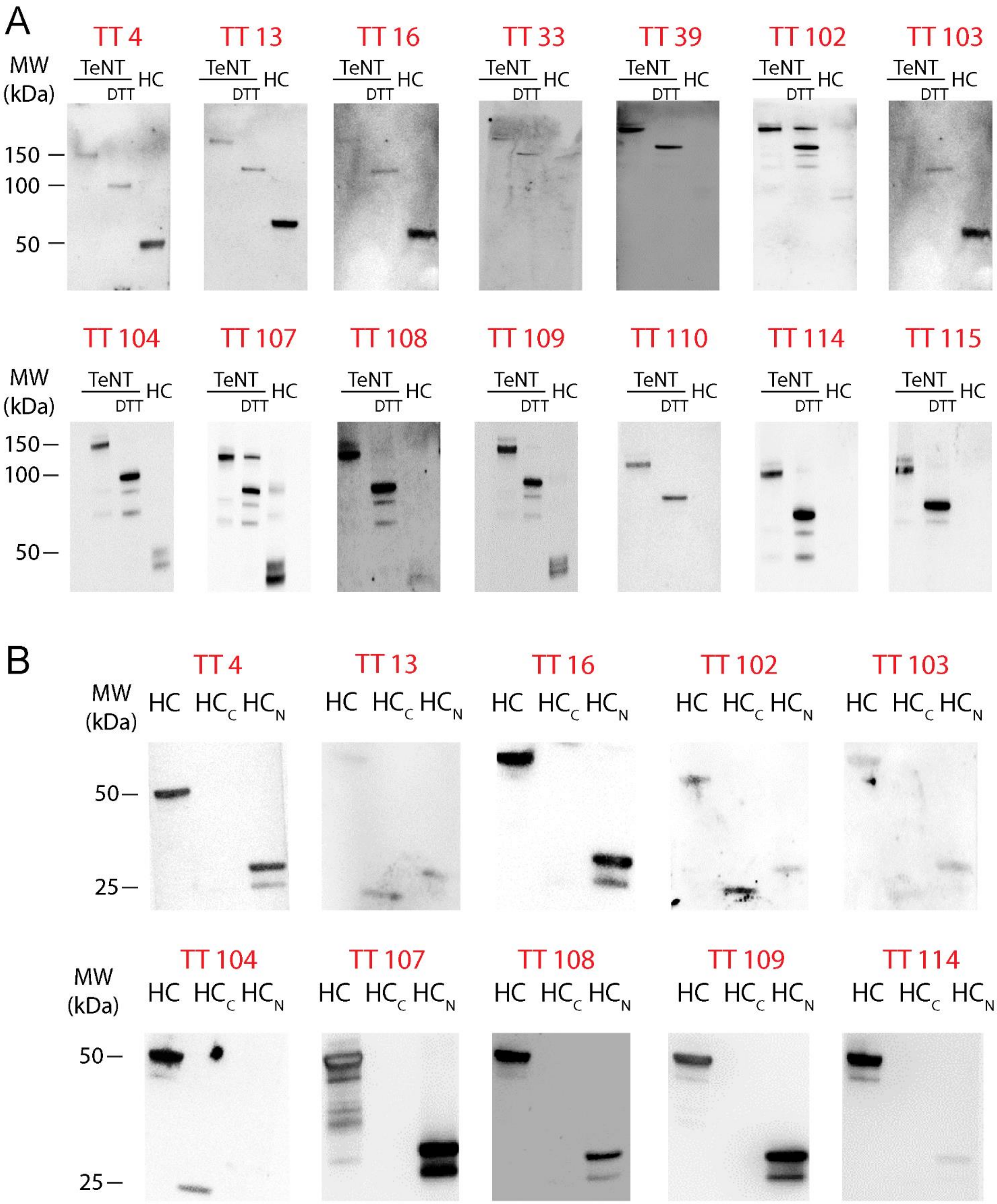

**Figure S2**: Structure of an immunoglobulin G showing its different domains and the site in the heavy chain where the stop codon has been inserted to generate the Fabs. Below are reported the amino acid sequences of the variable portion of the immunoglobulin G L chain (VL) and H chain (VH) of TT104-Fab and TT110-Fab.

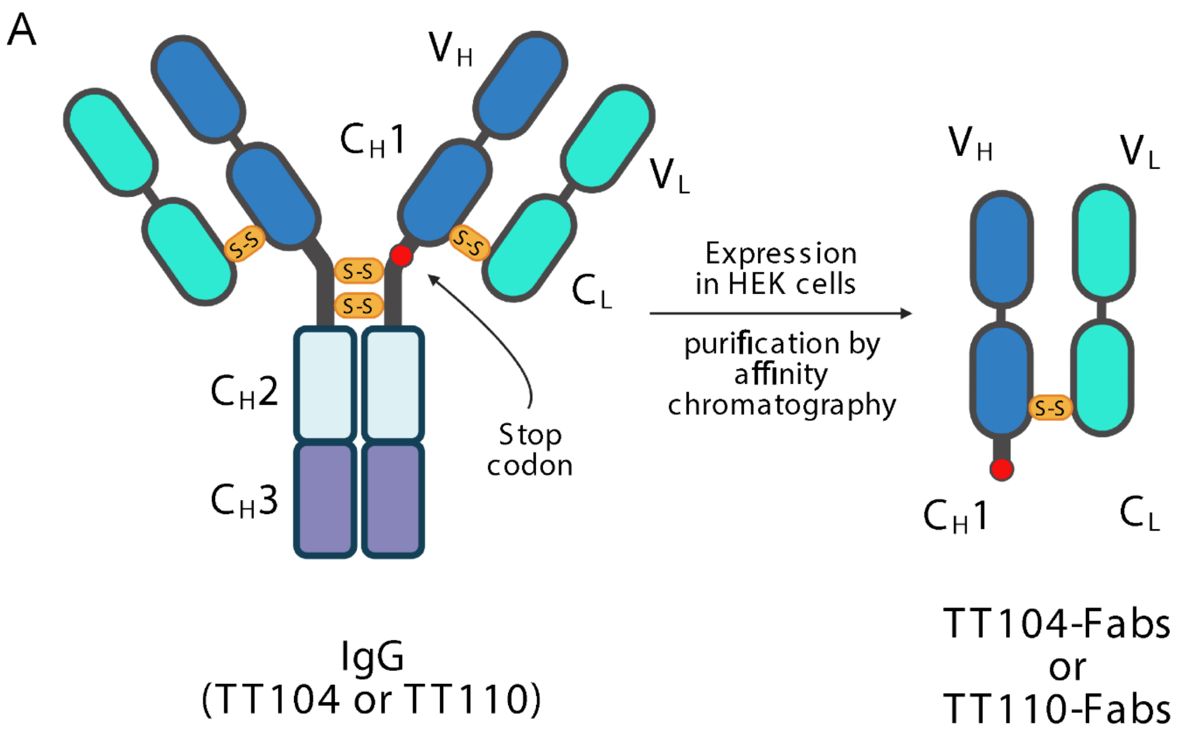

**Figure S3**: TeNT binding of TT104-Fab and TT110-Fab assayed by surface plasmon resonance. SPR analysis of the interaction between TeNT and TT104-Fab (A) and TT110-Fab (B). TeNT was immobilized on a TRIS-Ni-NTA sensor chip. Sensograms show the binding of incremental concentrations of Fabs on the sensor chip. The binding data were analyzed with a one site binding model.

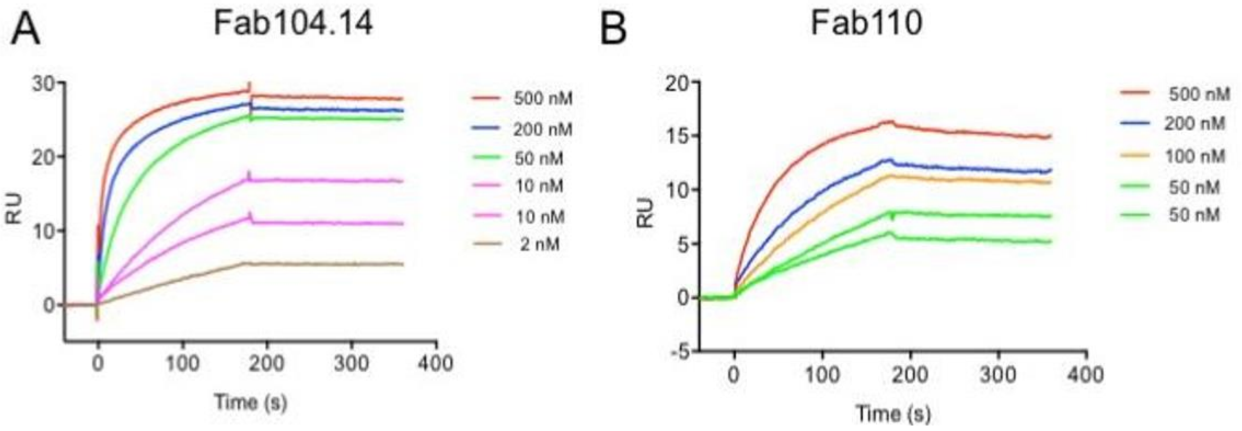


| | $K_a$ (1/Ms) | $K_d$ (1/s) | $K_D$ (M) |
|---|---|---|---|
| **TeNT-TT104-Fab** | $4,5*10^{-5}$ | $3,0*10^{-6}$ | $6,7*10^{-12}$ |
| **TeNT-TT110-Fab** | $6,3*10^{-4}$ | $2,5*10^{-4}$ | $3,0*10^{-9}$ |

**Figure S4**: Size exclusion chromatography (SEC) and native polyacrylamide electrophoresis analysis of TeNT immunocomplexes. TeNT (violet trace), TeNT and TT104-Fab (red trace), TeNT and TT110-Fab (grey trace) or TeNT and the two Fabs together (ternary immunocomplex, blue trace). SEC analysis was performed with a Superdex 200 10/300 column (GE Healthcare) and respective peaks were analyzed on native gels.

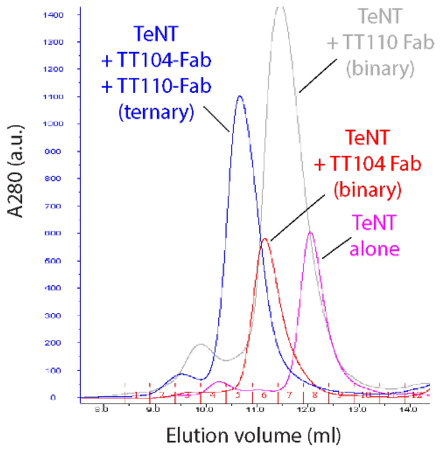

**Figure S5**: Cryo-EM image processing. A) from top to bottom: representative cryo-EM micrograph from untilted (left) and 30° tilted (right) data acquisition; selection of typical 2D class averages used for 3D reconstruction; results of the 3D global refinement; initial body used for multibody refinement; result of the multibody refinement and their associated FSC curves. B) contributions of all eigenvectors to the variance. The structure in fig. 3 C-D corresponds to the extreme of the first eigenvector. C) Histogram of the amplitudes along the first eigenvector shows a unimodal distribution.

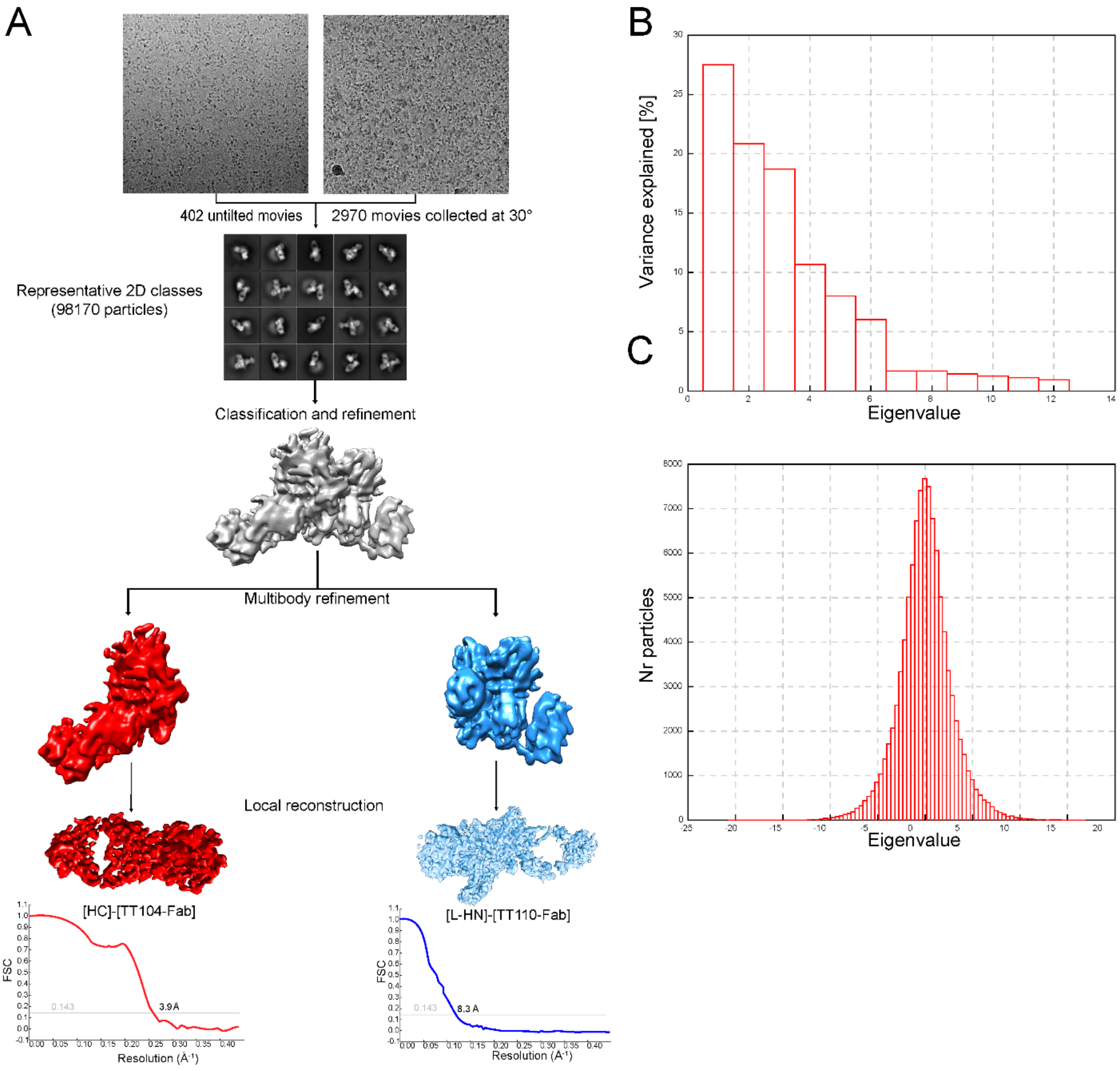

**Figure S6**: Representation of TT104-Fab bound to TeNT HC also in complex with nidogen. Nidogen binding is represented as previously modelled before (Bercsenyi et al., 2014). Spheres indicate the clashes predicted to occur between nidogen and TT104-Fab.

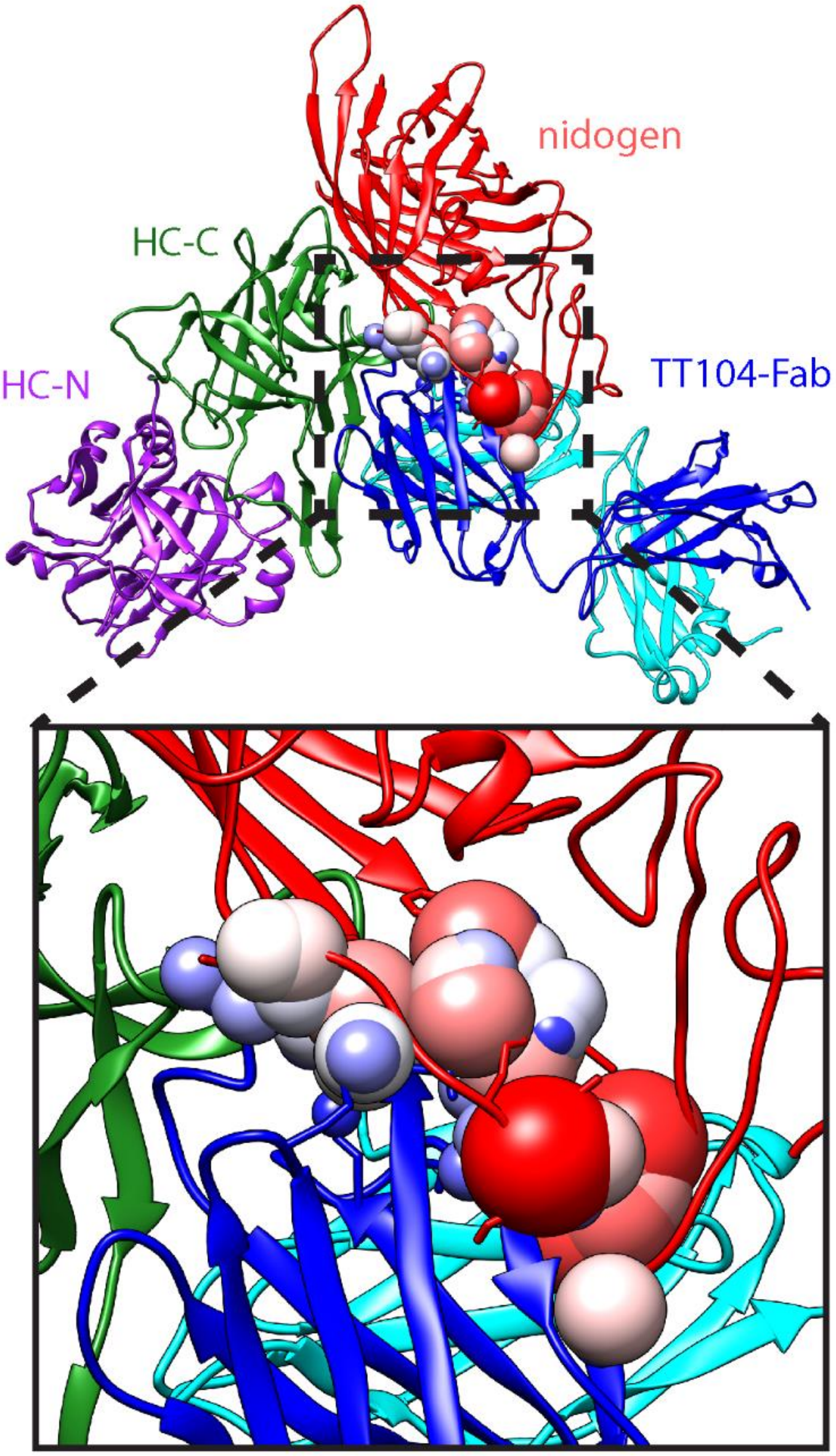

**Figure S7**. The BoNT switch is also present in TeNT. Representation of the HN domains of BoNT/A (left) and TeNT (right), where the BoNT switch is highlighted in red. Both structures display a very similar arrangement of this structural motif that was recently predicted to be the first part of the BoNT molecule that inserts into the membrane during the translocation process (Lam et al., 2018).

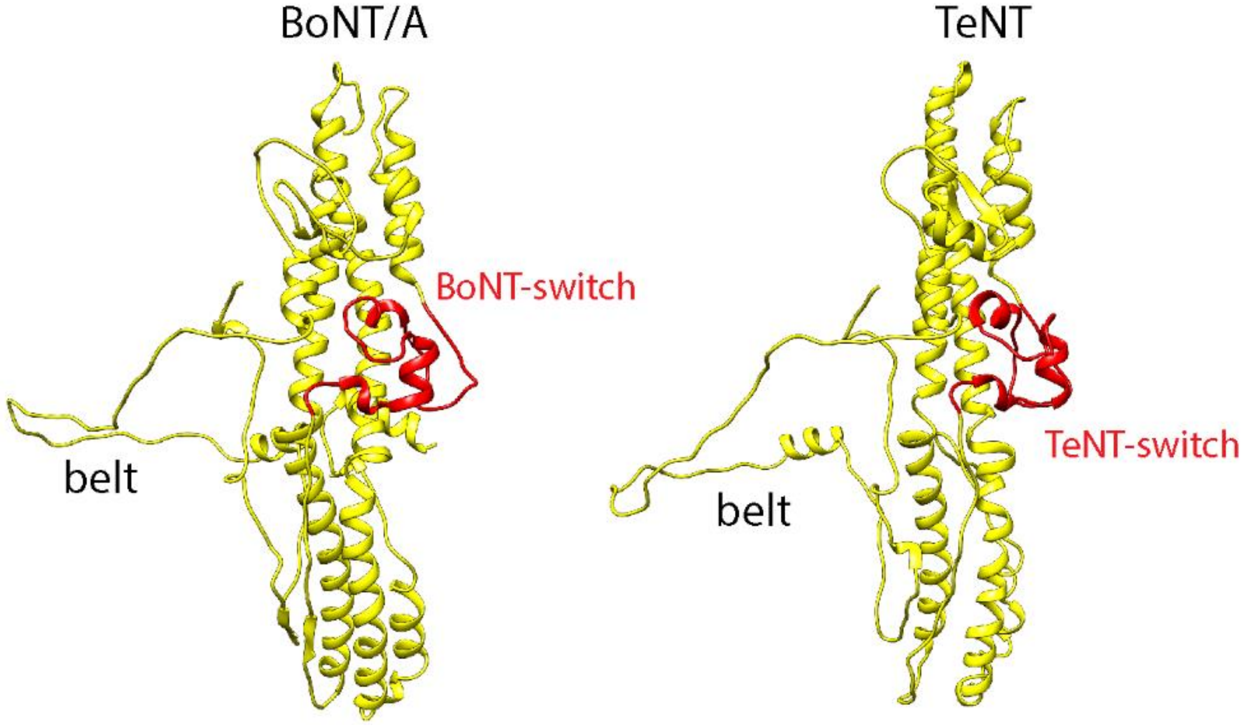

**Figure S8**: Cartoon representation of TT110-Fab bound to TeNT HN highlighting carboxylate residues possibly involved in the low pH induced structural change of BoNT and TeNT structure. The top panel shows the large area of the TeNT epitope recognized by the Fab derivative of the humAb TT110. The lower panel shows a close up view of the Fab interaction with the central helix of the "TeNT-switch", which corresponds to the $\alpha_B$ helix of the BoNT-switch in BoNT/A (Lam et al., 2018). The helices of the HN domain of TeNT, marked in red, correspond to the helices in $\alpha_A$ and $\alpha_C$ of the BoNT-switch, respectively. In orange it is highlighted another helix within the HN domain of TeNT which is part of the epitope recognized by the TT110-Fab. In green are the carboxylate residues suggested by the present study to play an essential role in the HN structural change caused by acidification inside synaptic vesicles.

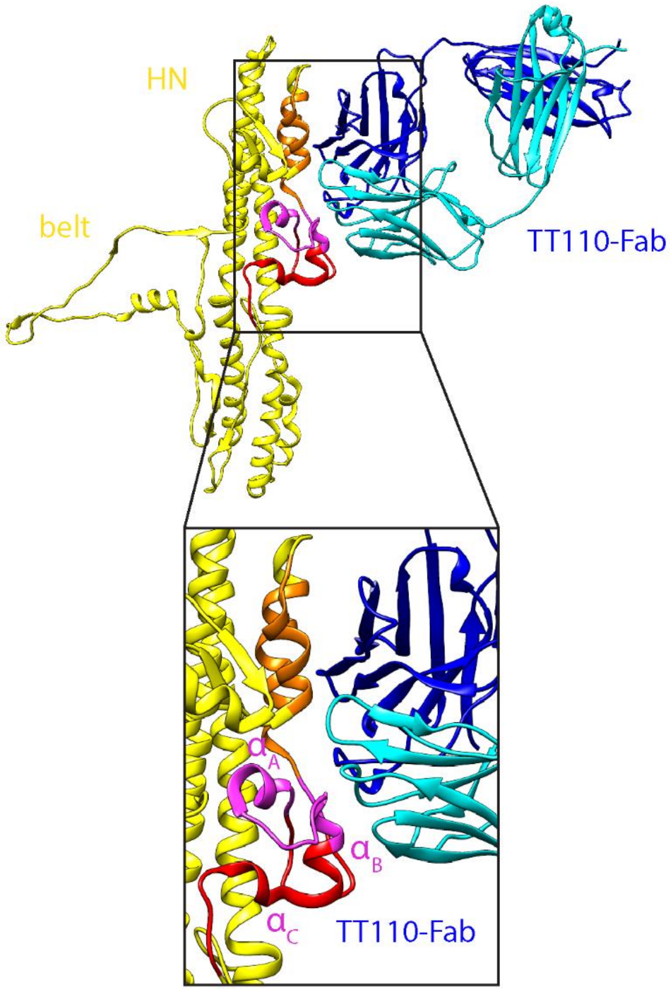

**Supplementary Table 1. Atomic details of TeNT-TT104-Fab interaction**. H-bond and salt bridge interactions and between the HC domain of TeNT and TT104-Fab, as detected by server PISA (https://www.ebi.ac.uk/pdbe/pisa/).

| | HC | Chain B | Chain A |
|---|---|---|---|
| H-bond | Lys1143 O | Ser32 OG | |
| H-bond | Gln1141 NE2 | Thr33 O | |
| H-bond | Asn1153 OD1 | Asn103 ND2 | |
| H-bond | Asn1153 ND2 | Asn103 OD1 | |
| H-bond | Tyr1202 OH | Ser31 O | |
| H-bond | Tyr1202 OH | Ser32 O | |
| H-bond | Tyr1150 OH | Asp102 OD1 | |
| H-bond | Asn1153 ND2 | | Tyr318 OH |
| H-bond | Ser1156 N | | Asn314 O |
| Salt Bridge | Arg1281 | Asp102 | |

**Supplementary Table 2. Model quality.**

| | HC- TT104-Fab | LC-HN-TT110-Fab |
|---|---|---|
| Residues/atoms | 870 / 6830 | |
| RMSD<br>bond length (Å)<br>Angles (°) | <br>0.009<br>1.16 | |
| MolProbity score | 3.43 | |
| Ramachandran plot (%)<br>Favored<br>Allowed<br>Outliers | <br>70.65<br>29.23<br>0.12 | |
| CC (mask/box) | 0.78 / 0.79 | |
